# Indirect environmental effects on the gut-brain axis in a wild mammal

**DOI:** 10.1101/2024.12.11.627940

**Authors:** Lauren Petrullo, Andrea Santangeli, Ralf Wistbacka, Arild Husby, Aura Raulo

## Abstract

Inconspicuous interactions between host physiological systems and resident microbial communities may underlie how animals respond to environmental change. For example, immunity and metabolism are regulated in part by the gut microbiota, which can be shaped indirectly by host neuroendocrine function via a “gut-brain axis”. Yet the sensitivity of this axis in wild vertebrates remains ambiguous. Here, we investigate covariation among environmental quality, glucocorticoids, and gut microbiota in a vulnerable population of Siberian flying squirrels (*Pteromys volans*) inhabiting a region impacted by variable rates of human disturbance. We test competing hypotheses related to direct versus indirect environmental effects (via the gut-brain axis) on adult and juvenile gut microbial communities. Adults housed a richer gut microbiota and had higher hair glucocorticoids that covaried with microbial composition, while juveniles lacked any hormone-microbiome covariation. Environmental quality (patch size, habitat diversity, and suitability) predicted variation in glucocorticoids but not variation in microbial diversity, suggesting no direct effects on gut microbiota. Instead, structural equation models revealed indirect environmental effects on microbiota via elevations in glucocorticoids in adults. Among juveniles, habitat-induced hormonal responses had no downstream effects on microbial diversity. Together, this provides evidence for age-dependent indirect environmental effects on gut microbial composition in a wild mammal by way of the host neuroendocrine system.

## INTRODUCTION

Anthropogenic change is rapidly reshaping ecosystems with profound effects on organismal fitness and population resilience (Sih et al., 2011). Underlying these effects are the molecular processes that can constrain or catalyze animal responses to both predictable and unpredictable environments (Fuller et al., 2010; Petrullo et al., 2025; Todgham & Stillman, 2013). Among vertebrates, the hypothalamic-pituitary-adrenal (HPA) axis is an evolutionarily conserved regulator of the physiological stress response, modulating host responses through a cascade of feedback interactions (Sapolsky et al., 2000). This cascade culminates in the production of glucocorticoid hormones (GCs), including cortisol (the primary GC in humans) and corticosterone [the primary GC in rodents, (Sapolsky et al., 2000)]. Acting as an integrator network, the HPA axis can transduce cues of changing conditions and coordinate phenotypic shifts that enhance fitness (Dantzer, 2023; Martin et al., 2011). Because the HPA axis is sensitive to a wide array of internal and external inputs, GC concentrations can index changes in host physiology following shifts in diet, predation risk, population density, temperature, and the social environment (Sheriff et al., 2011; Wingfield et al., 1998).

Notably, the HPA axis has both target and non-target effects on other physiological processes and organ systems. This includes interfacing with the symbiotic community of microorganisms that reside in the gastrointestinal tract (hereafter, gut microbiota), through a secondary “gut-brain axis” (Cryan & O’Mahoney, 2011). Through the gut-brain axis, neuroendocrine function may drive internal selective pressures that regulate within and among-host microbial variation (Cryan et al., 2019; Stothart et al., 2019). This variation can be further augmented by external drivers of microbial variation like diet, environment, and host-to-host microbial transmission (Baniel et al., 2021; Orkin et al., 2019; Raulo et al., 2021; Ren et al., 2017; Sarkar et al., 2024). In turn, these relationships may influence host fitness: gut microbiota can enhance pathogen protection (Spragge et al., 2023; Stecher & Hardt, 2011) and facilitate dietary flexibility (Cani et al., 2019; Kohl et al., 2016), but can also impair immune function and digestive efficiency (Pham & Lawley, 2014). Further, gut microbiota can themselves influence HPA axis function (Burokas et al., 2015), and consequently, host behavior (Davidson et al., 2020; Wu et al., 2021). The gut microbiota is thus governed by a series of complex internal feedbacks across the microbiota and the brain, generating covariation among HPA function, gut microbiota, and the broader environment with many causal linkages (**Figure 1**).

**Figure 1.**
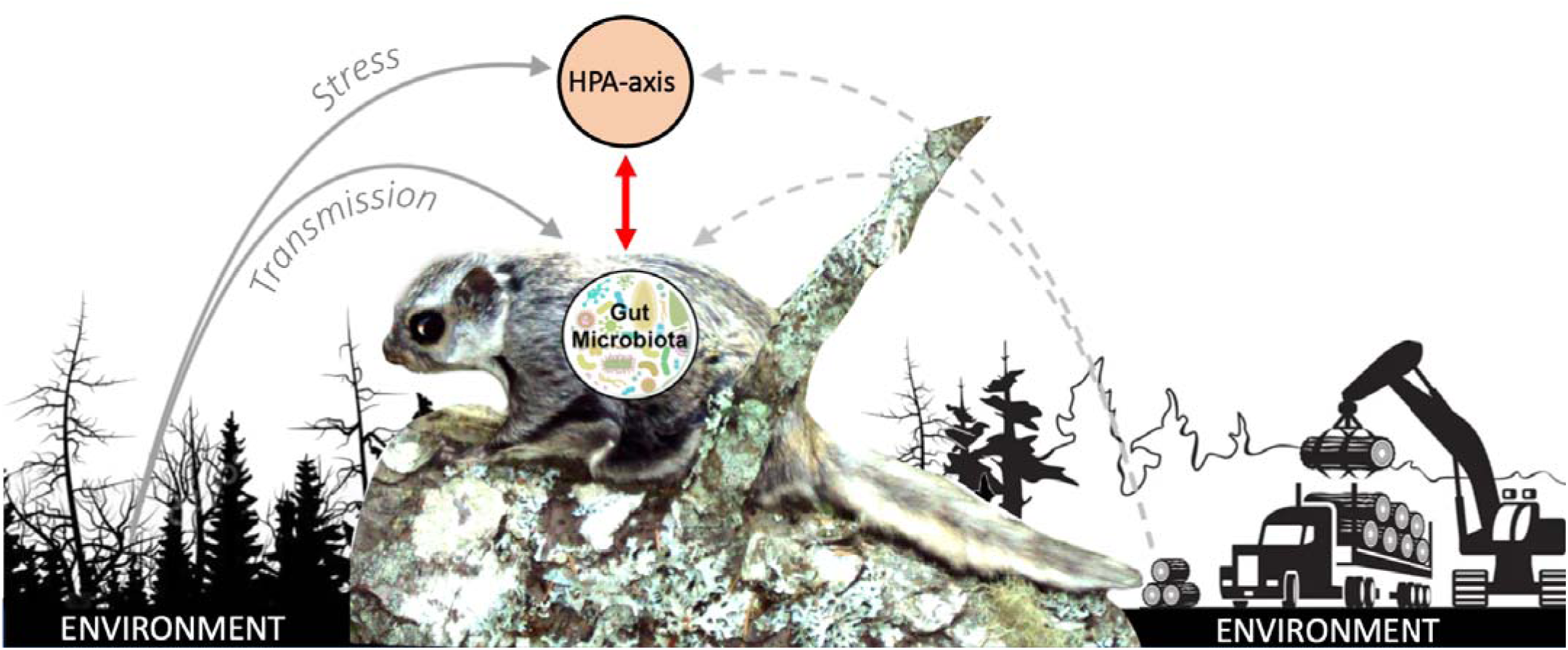
Causal linkages among the environment, gut microbiota, and hypothalamic-pituitary-adrenal (HPA) axis. The left and right side depict two different environments, one that is in a natural state and has low levels of disturbance and one that is structured by human action and has high levels of disturbance.

To date, empirical data and theory have largely supported the existence of a gut-brain axis in wild animals. For example, synthetic corticosterone administration reduces gut microbial diversity in yellow-legged gulls [*Larus michahellis,* (Noguera et al., 2018)]. Applications of these and similar findings [e.g., (Blanco et al., 2021)] remain limited, however, in the absence of ecologically-relevant, naturalistic stressors that may exert context-dependent effects on coupling of the neuroendocrine system and gut microbiota (Crespi et al., 2013). Studies in unmanipulated populations remain crucial to adding this context (Greyson-Gaito et al., 2020; Levin et al., 2016; Petrullo et al., 2022). Moreover, theoretical frameworks (e.g., the reactive scope model) suggest that inconspicuous feedbacks between an organism’s symbiotic microbiota, neuroendocrine system, and broader ecology impact host stress sensitivity (Houtz et al., 2022) and, in turn, ecological adaptation (Alberdi et al., 2016). The integration of environmental cues by the HPA axis could induce shifts in microbial community composition that promote reconfiguration of gut microbiota to anticipate or optimally track changing dietary landscapes (Baniel et al., 2021; Orkin et al., 2019). Alternatively, habitat degradation and habitat loss may increase GC production to adaptively mobilize energy stores (Busch & Hayward, 2009), but exert collateral effects on the microbiota by constraining taxonomic and/or functional diversity (Amato et al., 2013; Petrullo et al., 2022; Stothart et al., 2016).

In this study, we disentangle covariation across the HPA axis, gut microbiota, and environment in a wild small mammal. The Siberian flying squirrel (*Pteromys volans)* is an arboreal specialist species adapted to natural state old-growth boreal forests, and listed as vulnerable species within the European Union (Santangeli, Hanski, et al., 2013). Because their dietary niche is relatively narrow, they are particularly susceptible to changes in forest structure. Habitat loss due to commercial logging is a primary threat because it fragments their habitat into variably-sized patches of suitable, old-growth forest separated by unsuitable fields of young commercially-grown forest ((Santangeli, Wistbacka, et al., 2013; Wistbacka et al., 2018)). We test the hypothesis that habitat quality influences gut microbial variation in part through HPA axis activation (i.e., an indirect effect via the gut-brain axis). We take an integrative approach that includes structural equation modeling to test alternative hypotheses related to direct versus indirect environmental effects without requiring experimental manipulation of a vulnerable species (Eisenhauer et al., 2015; Grace et al., 2010; Laughlin & Grace, 2019). We expect that individuals inhabiting poorer quality habitats (smaller patch sizes, lower habitat diversity, and more unsuitable habitat) will exhibit higher GCs, indicating chronic HPA axis activation [(Sheriff et al., 2011), but see (Santangeli et al., 2019)]. In line with earlier studies, we predict that elevated GCs will alter gut microbial community ecology through microbial homogenization and a loss of microbial diversity (Noguera et al., 2018; Petrullo et al., 2022; Stothart et al., 2016, 2019).

## MATERIALS AND METHODS

### Study species and field data collection

Siberian flying squirrels (*Pteromys volans*) are arboreal rodents adapted to climax-state, spruce-dominated boreal forest. Within the European Union, most flying squirrels inhabit Southern Finland, which represents the western edge of their range across Eurasia. As old-growth forest specialists, flying squirrels are threatened by commercial forestry practices. Here, boreal forests are typically cut before they reach the climax state optimal for flying squirrels. As this is an area of intense commercial forestry, few disconnected patches of natural old-growth forest remain amidst younger commercial stands. Squirrel populations are thus scattered as a metapopulation across habitat patches of variable quality.

In June 2015, we sampled 78 Siberian flying squirrels (48 adults, 31 juveniles) from nest boxes across 69 sites in Finland. These sites are part of a long-term study on flying squirrels, and are deliberately positioned across a wide range of habitats [e.g., from urban park to natural old-growth boreal forests varying in their level of forestry-related disturbance, (Santangeli et al. 2013b; Selonen et al. 2014; Wistbacka et al. 2018]. We went through nest boxes in early June, at a time when flying squirrels are breeding and pups are on average ∼6 weeks old. We caught 79 individuals (48 adults and 31 pups). In most cases, nest boxes had either a solitary adult or one adult female and 1-4 pups. We collected data in situ, releasing squirrels to their point of capture <15 minutes within capture. Data collection included sexing and aging each individual, recording their body mass and femur length [see (Selonen et al. 2016) for more details], and tagging each individual with a subcutaneous PIT-tag.

### Hair and fecal sample collection

Hair is a minimally invasive sample medium that can index long-term stress exposure across weeks to months (Gormally & Romero, 2020). Hair GCs can thus capture chronic environmental stress more consistently than plasma GCs, which are affected by short-term stressors like handling at the time of collection (Desantis et al., 2016; Sheriff et al., 2011). We clipped a small amount (∼10 mg) of hair from the side of the tail of each individual squirrel to measure hair glucocorticoid content as a proxy for long-term stress exposure of our study subjects (Heimbürge et al., 2019). Hair sampling was done in a systematic and consistent way for all individuals: samples were cut as close as possible to the base of the hair and all samples were cut in the same exact way, from the same part of the tail, by the same person. This ensured that the GCs measured from the hair were as comparable as possible between individuals, as they would all relate to a similar time span of hair growth, during the same time of the year. Hair was stored in a dry place at room temperature before being sent to the Berlin Zoo.

Fecal samples were collected directly and opportunistically during animal handling and transferred into clean labeled polypropylene tubes with 1 ml of 98% ethanol. To avoid contamination, tubes and fecal pellets were handled with gloves, which were changed between every individual. Feces was stored in a fridge for the duration of the field work (2 weeks) and subsequently in a -80°C freezer, until they were shipped frozen to the Earth Microbiome Project at the University of Colorado for sequencing.

### Habitat quality measures

We used data from yearly habitat surveys processed in aGeographic Information System (GIS) to classify habitats across the study sites into three discrete classes following Wistbacka et al. (2018). Suitable habitat refers to breeding habitat for females, defined as layered spruce dominated mixed forests, containing trees of different size and age. Semi-suitable habitat is defined as monospecific forests (typically pine plantations) where flying squirrels can move but cannot breed because of lack of resources [e.g., food provided by deciduous trees or shelter provided by spruce trees, (Wistbacka et al., 2018)]. Unsuitable habitat is represented by open areas covered by young sapling stands or clear-cuts, roads, fields, water, or built up areas (i.e., areas unusable for flying squirrels to move, feed, and/or breed). We then calculated three measures of habitat quality: 1) patch size (area in ha), 2) patch diversity (different types of habitats present within 300 m of an individual’s nest box) and 3) proportion of suitable area within a 100 m radius from the center of each nesting site, which is where the nest boxes are located. The use of the 100m radius buffer is based on the known size of the core area used by breeding females, and was previously found the most relevant radius within which the habitat has the highest influence on the species (Santangeli, Hanski, et al., 2013; Wistbacka et al., 2018). The extended buffer of 300m used for patch diversity allows capturing more variance in this metric compared to a 100m buffer (which would generally have very low diversity and become nearly uninformative). While large, this 300 m buffer still captured the core area used by females during breeding, as well as extended areas, and habitats therein, in its surroundings. The amount of unsuitable habitat as well as size of the patch have been found to capture relevant habitat variation within the range typically used by flying squirrels (Hanski et al., 2000; Santangeli, Hanski, et al., 2013; Wistbacka et al., 2018).

### Measurement of hair glucocorticoids

Hair samples were washed with 100% methanol for 2 min. Methanol was decanted and the dried samples were powdered in a bead mill. Next, hairs were extracted with 400 μl 90% methanol and aliquots of 20 μl, diluted 1:2 with water, and analyzed in the enzyme immunoassay (EIA). Hydroxycorticosterone was quantified using a previously validated enzyme-linked immunosorbent assay (Voigt et al., 2004). Briefly, the assay used a polyclonal in-house antibody against cortisol-21-hemisuccinate (HS), coupled with bovine serum albumin (BSA) raised in rabbits, used together with the corresponding 21-HS-steroid coupled to peroxidase (HRP) as label for the EIA. Antibody cross-reactivity with different glucocorticoids was as follows: 4-pregnen-11a,17,21-triol-3,20-dione (cortisol) 100%, 4-pregnen-11ß,21-diol-3,20-dione (corticosterone) 13.2%, and 4-Pregnen-21-ol-3,20-dione (desoxycorticosterone) <□0.1%, respectively.

Standards were prepared in duplicates of 20 μl by 1:1 dilutions in assay buffer ranging from 0.2 to 100 pg/20 μl. All the duplicates, together with 100 μL cortisol-HRP conjugate in assay buffer (50 mM Na_2_HPO_4_/Na_2_HPO_4_, 0.15 M NaCl, 0.1% BSA, pH 7.4), were added to a plate, which was then incubated overnight at 4°C and then washed four times with washing solution. One hundred fifty microliter substrate solution was added per well (1.2 mM H_2_O_2_, 0.4 mM 3,3′,5, 5′-tetramethylbenzidine in 10 mM sodium acetate, pH 5.5), and the plates were incubated in the dark for 40 min. Fifty microliters of 4N H_2_SO_4_ were then added to stop the reaction and the samples were measured at 450 nm with a 12-channel microtiter plate reader (Infinite M 200; Tecan). Hormone concentrations were calculated using the Magellan software [Tecan; see (Finkenwirth et al., 2010)]. The sensitivity of the assay was defined as two standard deviations from the signal given by the zero blank and thus was 0.2 pg per well.

### Microbial DNA sequencing and bioinformatics

DNA extraction, amplification, and sequencing was performed in the Knight Laboratory at the University of Colorado in 2016, following Earth Microbiome Project protocols (see http://www.earthmicrobiome.org/emp-standard-protocols/16s/). In short, DNA extraction was done using MO Bio PowerSoil kits and PCR was run with primers 515F/806R, targeting the V4 region of the 16S rRNA bacterial gene (Caporaso et al., 2012). Resulting amplicons were sequenced using an Illumina MiSeq. Sequences were clustered into OTUs using the UPARSE pipeline (with de novo clustering using 97% similarity threshold and maximum expected error ≤2) and assigned taxonomy using GreenGenes (v XXXX). Further preprocessing was done with the *phyloseq* package in R (McMurdie & Holmes, 2013). Based on sample completeness curves produced with the iNext package (Hsieh et al., 2016), we chose to filter out samples with <10,000 reads as unrepresentative of the community diversity. Using the same package, we also derived asymptotic alpha-diversity estimates for each sample. Furthermore, we filtered all OTUs assigned to chloroplasts/Cyanobacteria. Finally, taxon abundances were normalized as proportions within their sample. Microbiome variation was then described through two set of metrics: richness (number of unique taxa) and Shannon diversity (an abundance-weighted measure of diversity) as measures of alpha diversity, and Jaccard similarity (proportion of shared taxa between two samples) as a measure of beta-diversity.

### Statistical analyses

#### Identifying drivers of variation in gut microbiota

To identify the factors that predicted variation in host gut microbiota, we performed a first marginal permutational analysis of variance (PERMANOVA) on microbiome beta-diversity (Jaccard similarity values) using the *adonis2* function in the *vegan* package (Oksanen et al., 2007). We first tested the effects of host age, sex, date, and family membership on microbiome composition in the full dataset (N=78). We then ran a second marginal PERMANOVA with family set as a blocking factor, controlling for age and sample date, to assess effects of the three habitat measures (habitat diversity, patch size, and amount of unsuitable area) on gut microbiota composition. We additionally tested for variation in homogeneity of dispersion as a function of host age using the *betadisper()* function and a generalized linear mixed-effects model (family = beta).

#### Differential abundance of microbial taxa

To identify the taxonomic differences between the gut microbial communities of adults and juveniles, we constructed a series of linear mixed effects-models using lme4 (Brooks et al., 2017). Models test how age (fixed factor) predicted the relative abundance of bacterial families (arcsine square root transformed), controlling for host sex, date of sample collection, and nest box/family (random effects). We accounted for multiple comparisons and an inflated false discovery rate for multiple hypothesis testing by adjusting all P values using the Benjamini-Hochberg correction (Benjamini & Hochberg, 2000).

#### Dyadic models

To test whether variation in glucocorticoids predicted variation in gut microbiota, we built dyadic Bayesian beta regression models using the *brms* package in R (R Core team, 2015), as previously validated (Raulo et al., 2021, 2024). We used pairwise Jaccard similarity values as our response variable and predicted it with pairwise measures for hair glucocorticoids, with sex similarity (0/1=same/different sex), age similarity (0/1=same/different age category), spatial distance (metric distance between nest boxes) and nest sharing (0/1=same/different nest) added as covariates. All covariates either naturally ranged from 0 to 1 or were scaled to do so, to make model estimates for all terms comparable. As Jaccard similarity is a proportional metric bound between 0 and 1, a logit link function was used. We chose to use Jaccard –which captures presence/absence rather than abundance–in this model over another beta-diversity measure because it better captures potential confounding effects of social microbial transmission (Raulo et al., 2021, 2024). As the modeled values are pairwise measures, and as such not independent of each other, a multi-membership random effect structure was built into the model to account for the effect of identity of both individuals in each pairwise comparison. The model was run first for all individual pairs in the data and subsequently separately for adults and pups, omitting age similarity as a covariate.

#### Effects of host traits and habitat quality on gut microbiota and glucocorticoids

We used linear mixed-effects models to examine gut microbial alpha-diversity (richness, Shannon) as a function of host traits and habitat quality. Shannon indices were tukey transformed to achieve normality of residuals. A first set of models included host age and sex as fixed covariates, and family membership as a random effect predicting gut microbial alpha-diversity with hair GCs and measures of habitat quality (patch size, patch diversity, and amount of unsuitable area). For models investigating the effects of habitat quality on gut microbiota, we included sex as a fixed factor, and sampling date and time as random factors given work suggesting diurnal patterns of GC secretion from circulation into hair (Sharpley et al., 2010, 2011).

#### Testing for direct and indirect effects using structural equation modeling

We constructed structural equation models (SEMs) to simultaneously test for direct and indirect effects of habitat quality on gut microbiota using *piecewiseSEM* (Lefcheck, 2016). Quantitative structural equation modeling is the gold standard for testing *a priori* causal hypotheses in complex ecological systems. It allows for the partitioning of direct and indirect (mediator) effects of multiple variables nested within one or more mixed-effects component models (Laughlin & Grace, 2019). The *piecewiseSEM* package specifically improves upon other more traditional SEM approaches because it accounts for smaller sample sizes through Akaike Information Criterion (AIC) model comparison and permits the inclusion of random effects and thus repeated sampling (Lefcheck, 2016). It has also previously been used to elucidate gut-brain axis connections in wild animals (Petrullo et al., 2022). First, we converted all categorical variables to numeric. To retain power, we built separate SEMs for each age class and tested only the measures of habitat quality and covariates that were found to significantly predict GCs and microbial diversity in our previous models. For each SEM, the first component model included hair GCs (log-transformed) as the dependent variable, habitat quality (patch size, diversity or amount of unsuitable habitat) and sex as fixed effects, and family membership as a random effect. The second component model included gut microbial alpha-diversity (richness or Shannon diversity) as the dependent variable, hair GCs as the fixed effect and family membership as a random effect. Component models were compiled into one and structural equation models were run using the *psem()* function. Model fit was assessed using AIC(c), and standardized coefficients were extracted to allow for comparison of model estimates across the component models (Lefcheck, 2016).

## RESULTS

### 1. Intrinsic and extrinsic drivers of gut microbial variation in flying squirrels

The Siberian flying squirrel gut microbiota was composed of bacteria from across 14 phyla, with most microbiota from Firmicutes, Bacteroidetes, Actinobacteria, and Proteobacteria (**Figure 2A)**. Within these phyla, samples contained microbiota from 97 families (**<u>Figure S1</u>)**. Specifically, *Lachnospiraceae* (mean relative abundance = 39.4% across samples), *Ruminococcaceae* (21%), and *Bacteroidaceae* (20.4%) were the most highly abundant families, with all other families contributing less than 5% to the total gut microbial community (**<u>Figure S1</u>)**.

**Figure 2.**
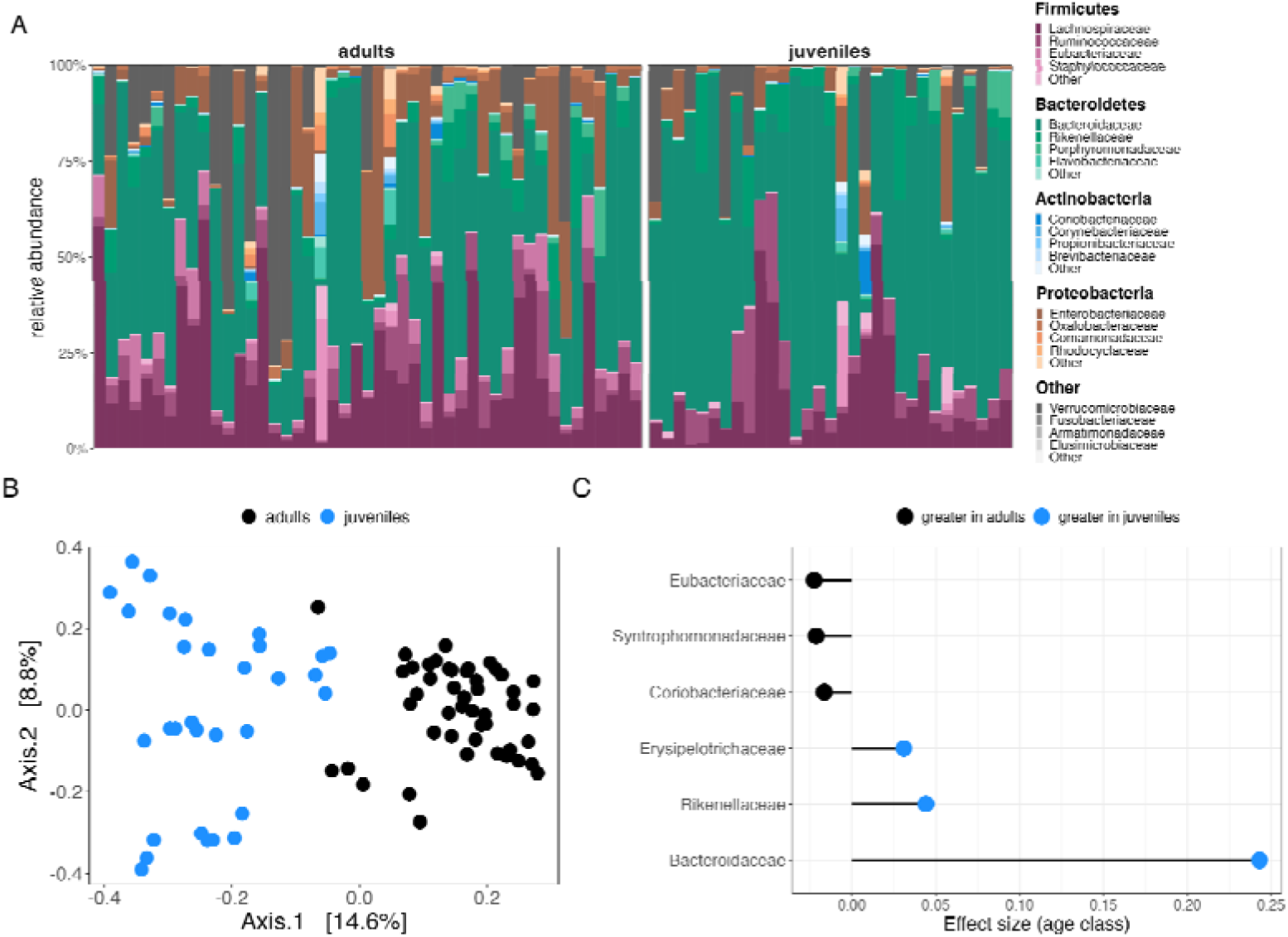
Age effects on microbial community composition. (**A)** The flying squirrel gut microbiota was dominated by microbiota from four main phyla. The most abundant microbial families (ranked) are shown below each phyla in the legend. **(B)** Principal coordinates analysis of Jaccard similarity across all fecal samples (N =78) by age class. **C)** Differential relative abundances of bacterial families (P_FDR_ > 0.05) as a function of age class. Lollipop plot depicts effect sizes from linear mixed-effects models testing the differential relative abundance of bacterial families (relative abundance > 0.05%) as a function of age class.

A marginal PERMANOVA on Jaccard similarity revealed that an individual’s nest box, which reflects family membership, explained a substantial amount of variation in gut microbiota composition (*R*^2^ = 0.47, F = 1.17, *p* = 0.001). Additionally, host age additionally explained some of the variance (*R*^2^ = 0.03, F = 1.89, p = 0.001, **<u>Table S1</u>, Figure 2B).** There was no effect of host sex on gut microbial composition (**<u>Table S1</u>**). Compared to adults, juvenile squirrels exhibited significantly greater compositional variability (greater distances to group centroid in Jaccard compositional space; estimate ± SE: 0.14 ± 0.03, z = 4.84, *p* < 0.001, **<u>Figure S2</u>, <u>Table S3</u>**), demonstrating greater inter-individual differences in microbial composition during early development compared to later life. At the taxonomic level, juveniles housed significantly higher relative abundances of gut microbiota from the families *Bacteroidaceae*, *Rikenellaceae*, and *Erysipelotrichaceae* (**Figure 2C**, **<u>Table S4</u>**). By contrast, adults housed greater relative abundances of *Coriobacteriaceae, Syntrophomonadaceae, and Eubacteriaceae* **(Figure 2C, <u>Table S4</u>).**

### 2. Glucocorticoids predict variation in gut microbial diversity and similarity

For individuals from which we had hair GCs (N=43), we found that juveniles exhibited substantially lower GCs than adults (linear mixed-effects model predicting hair GCs, estimate ± SE: -0.79 ± 0.13, t = -6.24, *p* < 0.0001, **<u>Table S5</u>, <u>Figure S3</u>)** as well as lower gut microbial alpha-diversity (linear mixed-effects model predicting microbial richness (number of OTUs), estimate ± SE: -327.81 ± 36.99, t = -8.86, *p* < 0.0001; predicting Shannon Index, -40.87 ± 4.76, t = -8.59, *p* < 0.0001; **<u>Table S6</u>, <u>Figure S3</u>**). Covariation between GCs and microbial diversity was also age-dependent: among adults, higher GCs predicted greater gut microbial Shannon diversity (linear mixed-effects model, estimate ± SE: 0.48 ± 0.02, t = 28.61, *p* < 0.0001), with a marginal effect on microbial richness (estimate ± SE: 26.82 ± 14.02, t = 1.91, *p* = 0.07, **<u>Table S7</u>, Figure 3A**). However, there was no relationship between GCs and either measure of microbial alpha-diversity among juveniles (**<u>Table S7</u>, Figure 3B**).

**Figure 3.**
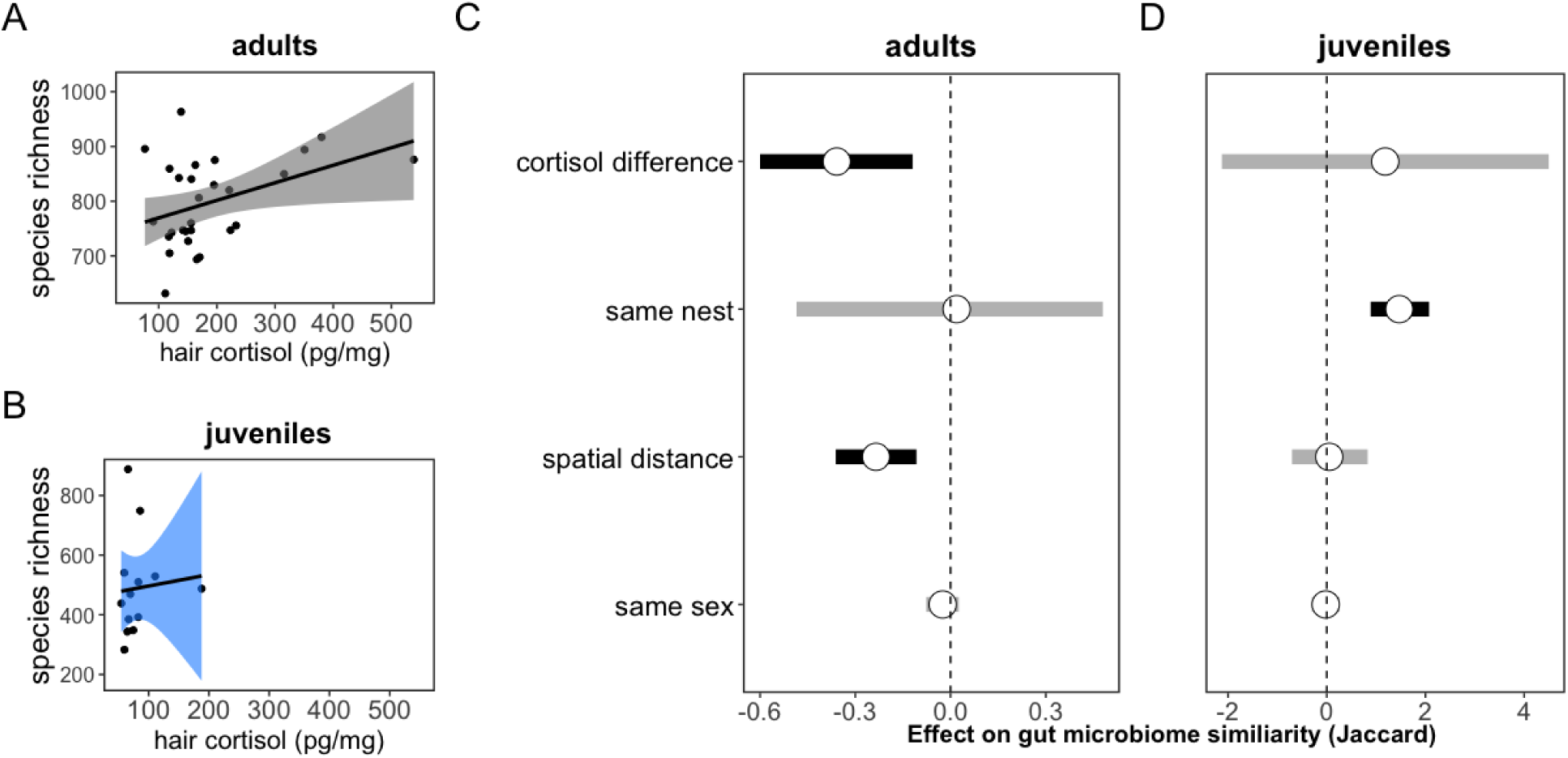
Covariation between glucocorticoids and gut microbiota is present in adults but absent in juveniles. **(A)** Adult GCs were positively associated with microbial alpha-diversity (species richness). **(B)** Among juveniles, there was no relationship between GCs and microbial alpha-diversity. **(C)** Adult squirrels with less similar hair GC concentrations and larger spatial distances between them had less similar microbiota (i.e., effect on the negative side, credible intervals do not overlap 0). **(D)** Among juveniles, differences in GCs did not predict gut microbiome similarity but juveniles residing in the same nests shared significantly more gut microbial taxa than juveniles residing in separate nests. Posterior means (points) and their 95% credible intervals (lines) are plotted from dyadic Bayesian GLMM with pairwise microbiota similarity among squirrels (Jaccard similarity) as the response.

To further parse the relationship between GCs and gut microbiota, we modeled microbial beta-diversity (Jaccard similarity) as a function of similarity in GCs, controlling for spatial distance, sex, and nest sharing (Raulo et al., 2021, 2024). Adult squirrels with more similar GC concentrations shared a greater proportion of microbial taxa than those with more divergent GC concentrations (dyadic Bayesian generalized linear-mixed effects model, posterior mean = - 0.36, CI = -0.60 to -0.12; **Figure 3C, <u>Table S8</u>**). Additionally, adults in closer spatial proximity to one another also had more similar gut microbiota (posterior mean = -0.23, CI = -0.36 to -0.11; **Figure 3C, <u>Table S8</u>**). By contrast, among juveniles, neither spatial distance nor differences in GC concentrations predicted gut microbiome similarity (<u>Table S6</u>). Instead, similarity among juvenile microbial communities similarity was predicted only by nest box, such that juveniles shared a greater proportion of their gut microbial taxa with their nest-mates (i.e., siblings, posterior mean = 1.47, CI = 0.89 to 2.07; **Figure 3D, <u>Table S9</u>**). Sex (same versus different sexes) did not predict microbial similarity in either age class.

### 3. Age-dependent environmental effects on the HPA axis, but not microbiota

Habitat quality was associated with variation in hair GCs, but the magnitude and direction varied as a function of age class and habitat measure. Adults inhabiting smaller patches had higher GCs (estimate ± SE: -0.15 ± -0.07, t = -2.16, P = 0.038), with the opposite trend in juveniles (**Figure 4A, <u>Table S10</u>**). Habitat diversity positively predicted GCs in juveniles (estimate ± SE: 0.34 ± 0.14, t = 2.34, P = 0.026) but not in adults (**Figure 4B, <u>Table S10</u>**). The amount of suitable habitat had no effect on GCs in adults or juveniles (**Figure 4C, <u>Table S10</u>**). Moreover, none of the three habitat measures had direct effects on gut microbial diversity (richness or abundance-weighted Shannon diversity) in either age class (**<u>Table S11</u>**).

**Figure 4.**
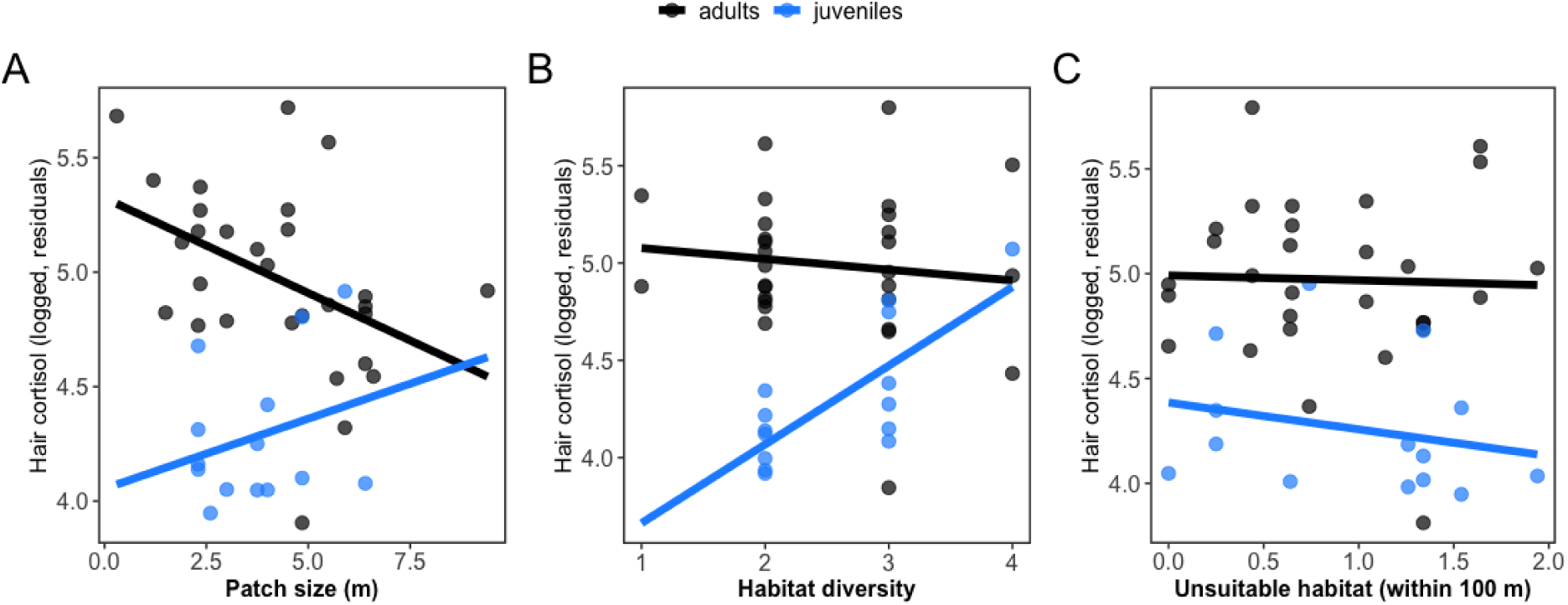
Adults and juveniles exhibit differential hormonal sensitivity to measures of environmental quality. **(A)** Patch size and **(B)** habitat diversity explain variation in GCs in an age-dependent manner, with no effects of **(C)** amount of unsuitable habitat on GCs in either adults or juveniles. Scatterplots show results from linear mixed-effects models. Points reflect samples (partial residuals), lines indicate regression lines from generalized linear-mixed effects models.

### 4. Environmental quality shapes gut microbiota indirectly via glucocorticoids

Because habitat quality predicted GCs, and GCs had a positive effect on gut microbial richness among adults, we used structural equation modeling to test the hypothesis that habitat-induced variation in GCs mediated downstream effects on gut microbial diversity. Among adults, smaller patches increased GCs (standardized β = -0.39, *P* = 0.15), which in turn increased microbial diversity (standardized β = 0.39, *P* = 0.46, **Figure 5A, <u>Table S12</u>**). Among juveniles, there was no relationship between GCs and microbial diversity. However, a more diverse habitat increased GCs (standardized β = 0.84, *P* = 0.001), but had no subsequent effects on microbial diversity (**<u>Table S12</u>, Figure 5B**).

**Figure 5.**
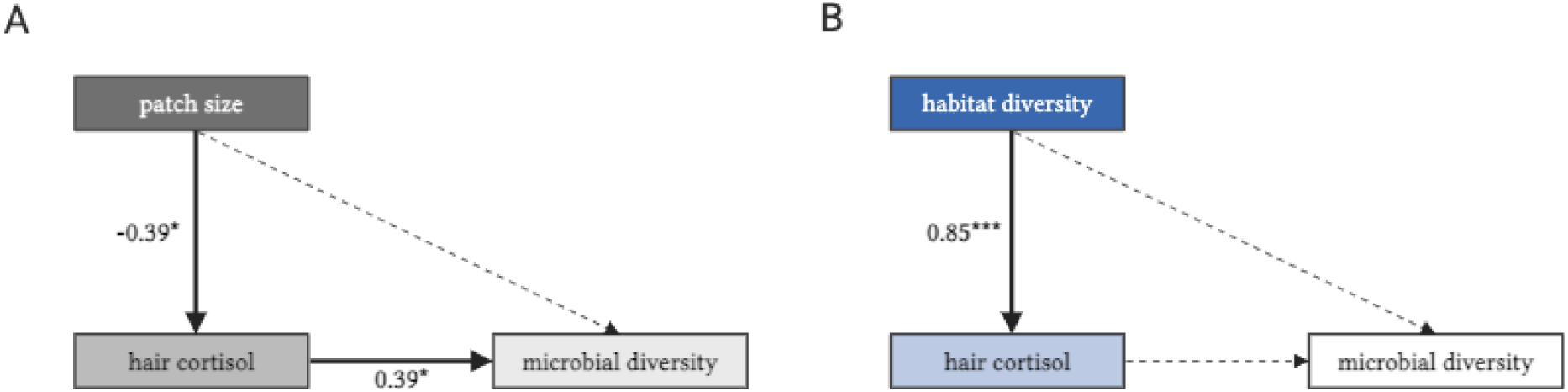
Age-dependent and indirect effects of habitat quality on gut microbiota. Among adults, **(A)** smaller patch sizes increased richness via elevations in GCs. Among juveniles, **(B)** elevations in GCs induced by habitat diversity had no subsequent effect on richness in pups. Solid arrows indicate significant paths, dashed arrows indicate non-significant paths. Covariates not shown for visualization purposes (see Table S12 for details). Model estimates for significant paths are shown with significance level (\**P* < 0.05, ** P < 0.01, \*\*\**P* < 0.001) and reflect standardized beta weights from mixed-effects component models controlling for sex, sample date, time, and nest box.

## DISCUSSION

Environmental change can activate both adaptive and maladaptive physiological responses associated with the secretion of glucocorticoid hormones by the HPA axis (Dantzer et al., 2014; Jaimez et al., 2012; Romero, 2004; Sheriff et al., 2011). Here, we show that flying squirrels responded hormonally to variation in habitat size, such that hair GCs were higher among squirrels inhabiting smaller patches. Many arboreal animals, including tree squirrels and flying squirrels, are adapted to mature forest habitat (Selonen & Mäkeläinen, 2017). Changes in forest structure that reduce the size of usable habitat can lead to fragmentation and resource scarcity that elevates GCs [small mammals: (Boyle et al., 2021), nonhuman primates: (Carlitz et al., 2016; Rangel-Negrín et al., 2009)]. Smaller patches can also increase proximity to external stressors (e.g., humans, predators) at habitat edges, increasing GC production [e.g., squirrel gliders, (Brearley et al., 2010, 2012)]. Elevated GCs among squirrels inhabiting smaller patches may thus reflect psychosocial stress, increased vigilance, and/or anti-predator behaviors (Creel et al., 2009; Voellmy et al., 2014). Moreover, when habitats shrink, squirrels may mobilize energy stores to cope with the increased metabolic burden associated with traveling greater distances for food or mates (Crespi & Denver, 2005). These responses may lead to chronic HPA axis activation, causing increased accumulation of GCs into hair (Russell et al., 2012).

Unlike environmental effects on the HPA axis, we found no evidence for direct effects of any measure of habitat quality on gut microbiota. Ambient temperature and vegetation explain gut microbial variation with longitudinal sampling in similar populations, suggesting our cross-sectional analyses may have been temporally constrained (Liu et al., 2019). Nonetheless, we found support for indirect, age-dependent environmental effects that run through the HPA axis. Unlike the reduction in microbial diversity documented in other wild animals [e.g., (Petrullo et al., 2022; Stothart et al., 2016)], adult squirrels with higher GCs housed a richer gut microbiota with greater abundance-weighted alpha-diversity. At the highest level, GCs may mechanistically enhance microbial diversity via stress-related behaviors like social grooming in highly social species like red-bellied lemurs ((Raulo et al., 2018)). This is an unlikely explanation for microbial diversification in less social systems like Siberian flying squirrels, though behavioral changes mediated by GCs may still play a role: higher GCs could promote increased foraging or vigilance, which may increase opportunities for social and environmental microbial transmission (Voellmy et al., 2014). At the molecular level, microbial diversification may occur via bacterial sensitivity to GC-induced changes in the biochemical environment of the gastrointestinal tract (e.g., pH, barrier function), which can “fight” or “feed” different microbiota (Karl et al., 2018). The immunomodulatory capacity of GCs may diversify microbiota through selective targeting of gut bacteria by the mucosal immune system, namely via secretory immunoglobulin A (sIgA). A change in microbial targeting by sIgA could reduce dominance of “core” microbiota and open niche space for rarer and less abundant bacteria, increasing alpha-diversity (Catanzaro et al., 2019). Such sIgA-mediated effects may be further amplified seasonally as host diets change and microbial diversification is needed to maintain the metabolic flexibility that facilitates dietary transitions, or as metabolic rates change to meet thermoregulatory demands (Han et al., 2023).

While a diverse microbiota is typically viewed as beneficial for hosts, diversity does not consistently index better host health or fitness (Williams et al., 2024). Diversification resulting from elevated GCs could reflect a loss of functional homogeneity generated by canalized microbial responses to ecological cues (e.g., specialized diets). If host HPA axis activation acts as an internal perturbation, it could cause microbial “scatter” that increases alpha-diversity, similar to the Anna Karenina principle (Zaneveld et al., 2017). A diversified microbiota could thus have potentially deleterious effects for hosts if it interrupts the adaptive calibration of microbial communities in response to internal or external cues (Petrullo et al., 2025). Yet, adult squirrels had more similar microbiota (i.e., lower beta-diversity) if their GC concentrations were also more similar, suggesting that microbial communities do not vary idiosyncratically in response to HPA axis activation. Instead, microbial diversification in the footsteps of HPA axis activation may reflect a coordinated suite of adaptive physiological changes to stress. Nonetheless, the directionality and magnitude of these effects is likely to vary across species, resulting in apparent discrepancies in GC-induced enhancement or depletion of microbial community diversity (Davidson et al., 2020).

Our earlier work found no relationship between habitat quality and GCs, but we did not test for age-dependence in these patterns (Santangeli et al., 2019). Adult flying squirrels interact with more dimensions of their environment more directly than juveniles, which are largely bound to their nests and exposed to local phenomena and sibling competition. We found that juveniles exhibited divergent hormonal sensitivity to environmental conditions relative to adults: those inhabiting more diverse habitats exhibited higher GCs, and this was the only environmental variable predictive of the HPA axis during development. A diverse habitat is typically viewed as an indicator of a healthy ecosystem (Rapport, 1989), and this could translate into higher GCs if juveniles are exposed to a greater diversity of predators, competitors, or nest parasites. Overall, that the hormonal response was divergent between adults and juveniles suggests differential sensitivity to environmental stress as a function of life history stage (Crespi et al., 2013). In line with this interpretation, while habitat disturbance increases host glucocorticoids in adult mammals [e.g., reviewed in (Dantzer et al., 2014)], this effect becomes inconsistent when juveniles are included in such analyses (Pérez-Ortega & Hendry, 2023).

Because both gut microbiota and the HPA axis exhibit independent and age-dependent sensitivity to intrinsic and extrinsic inputs (de la Cuesta-Zuluaga et al., 2019; Sapolsky & Meaney, 1986), it is perhaps unsurprising that their coordinated function as an integrated axis also varied with host age (Scott et al., 2017). Indeed, juveniles housed a substantially different gut microbiota than adults, enriched in bacterial families involved in the processing of high-fat diets like milk during early life in mammals [e.g., *Bacteroidaceae, Rikenellaceae, (Comstock, 2009)*]. Juveniles also exhibited lower GCs than adults. Because HPA axis responses are typically attenuated during early development in mammals (Sapolsky & Meaney, 1986), GC increases may not be robust enough to influence microbial composition at the community level in juveniles (i.e., dose-dependency). Unlike adults, juveniles are also continually exposed to conspecific microbiota in the nest: we found that nest box was the only significant predictor of juvenile pairwise microbial similarity. Social microbial transmission across littermates may eclipse glucocorticoid-induced differences in gut microbiota across individuals through a swamping effect, particularly if the pathways linking microbiota to the host endocrine system have not yet been sufficiently primed (Burokas et al., 2015). Further, maternal transfer of antibodies like sIgA to juveniles via milk may block or offset microbial change mediated by the HPA axis by augmenting endogenous juvenile sIgA production in the gut (Ding et al., 2021). Finally, GC accumulation in the hair shaft is an aggregate measure of host GC production over an extended period of time. It may thus be too coarse a measure to capture brief but pronounced hormonal responses if these responses are more variable in juveniles than in adults (Gormally & Romero, 2020). Ultimately, because developmental decoupling of the HPA axis and gut microbiota during early life may explain, at least in part, age-related differences in gut microbiota (Heitlinger et al., 2017), we encourage future gut-brain axis research to consider host life history stage.

Nevertheless, as interest in how the gut-brain axis can inform animal ecology and evolution grows, so do the number of outstanding questions. For example, which dimensions of host reproduction and longevity are impacted by microbial responses to ecologically-triggered hormonal responses? Structural equation modeling offers one approach for capturing gut-brain axis relationships in wild, vulnerable populations in which experimental manipulations are unethical or impossible (Lane, 2006). However, it cannot account for the bidirectionality inherent to the gut-brain axis (Burokas et al., 2015), as gut microbiota can themselves produce hormones and neurotransmitters that influence central nervous system function (Carabotti et al., 2015). Controlled field-based experiments and ‘omics approaches remain necessary next steps to isolate the molecular pathways linking gut microbiota to the brain within an ecological context (Davidson et al., 2020; Greyson-Gaito et al., 2020).

## ACKNOWLEDGEMENTS

We thank Se Jin Song and the Earth Microbiome Project for sequencing the fecal samples used in this work as part of their species catalog of gut microbiomes. We further give thanks to Sarah Knowles for support with field equipment, Ben Dantzer for help with early conceptualisation of this research idea, and the online forum Animal Microbiome Research Group for bringing the first and last authors together. Sample collection was supported by funding to RW from Svensk-Österbottniska samfundet, Societas Pro Fauna et flora Fennica, Vuokon Luonnonsuojelusäätiö, and Suomen Luonnonsuojelun säätiö. We also thank the National Science Foundation (DEB-2010726 to LP) and the University of Arizona.

## DATA AVAILABILITY STATEMENT

All data and code for this project are available at the following figshare repository: https://figshare.com/s/fdb14e7ca9ca6f269f42 (reserved DOI: 10.6084/m9.figshare.27616236 to at time of publication). An additional tutorial on the Bayesian dyadic multi-membership models can be found at AR’s github page: https://github.com/nuorenarra. All raw sequence data are publicly available at the NCBI SRA under PRJNA1283584.

## SUPPLEMENTAL MATERIAL

**Table S1.**
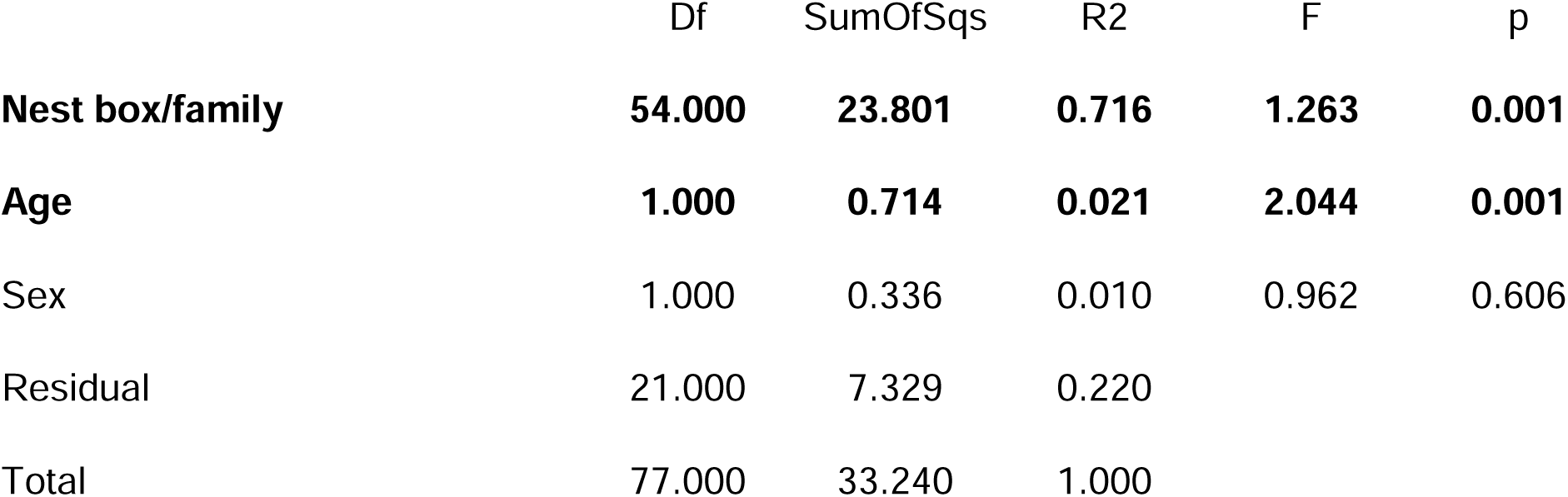
Intrinsic predictors of gut microbial variation. Results from a marginal PERMANOVA testing effects host age, sex, and family membership/nest box on Jaccard similarity, with 999 permutations.

**Table S2.**
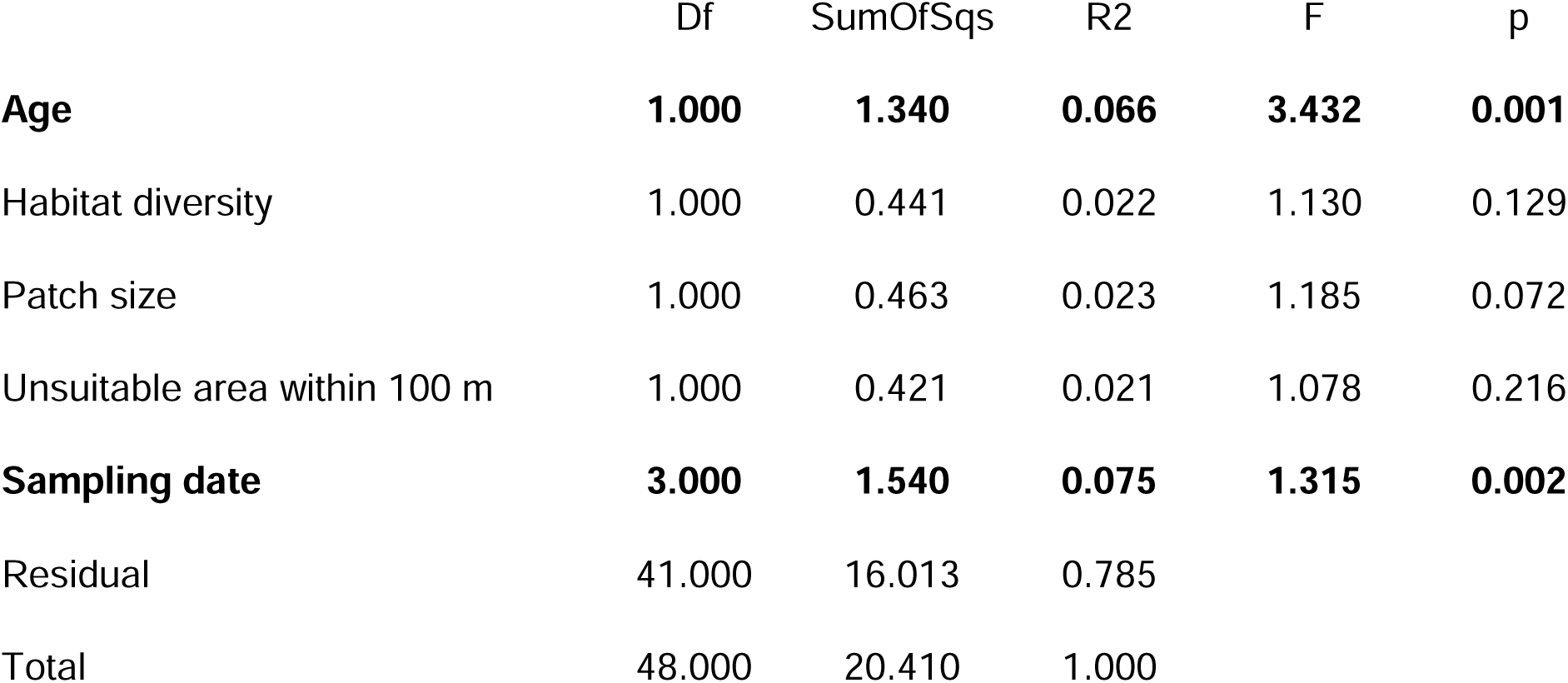
Extrinsic and methodological predictors of gut microbial variation. Results from a marginal PERMANOVA testing effects of habitat quality (habitat diversity, patch size, and amount of unsuitable area within 100 m of an individual’s nest box) and date of sample collection on gut microbial composition (Jaccard similarity), controlling for nest box/family membership (blocking factor).

**Table S3.**
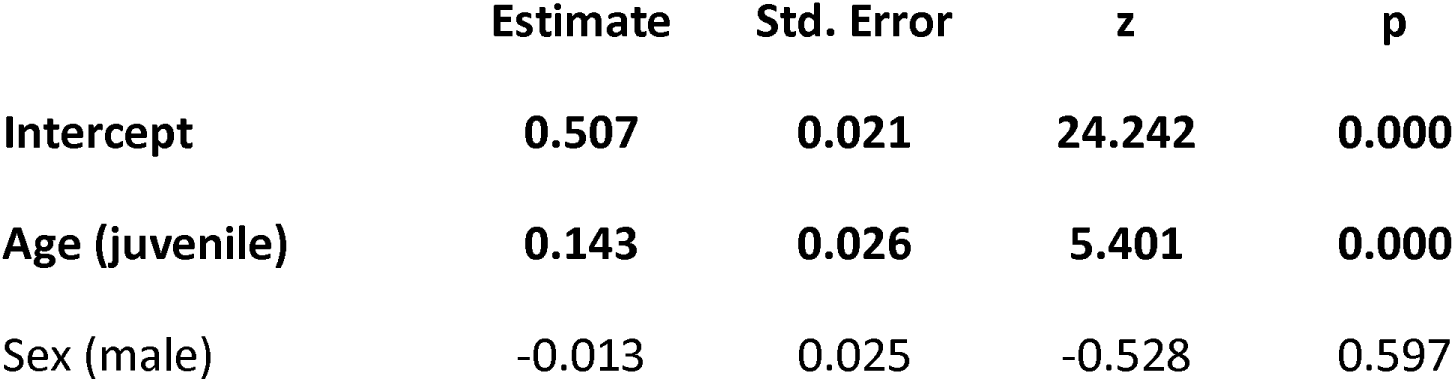
Juvenile flying squirrels exhibit greater inter-individual variability in gut microbial composition than adults. Results from generalized linear mixed-effects model (family = beta) testing the effects of age class on distance to group centroid in a Jaccard similarity compositional space. Model controlled for sampling date, family membership/nest box, and host sex.

**Table S4.**
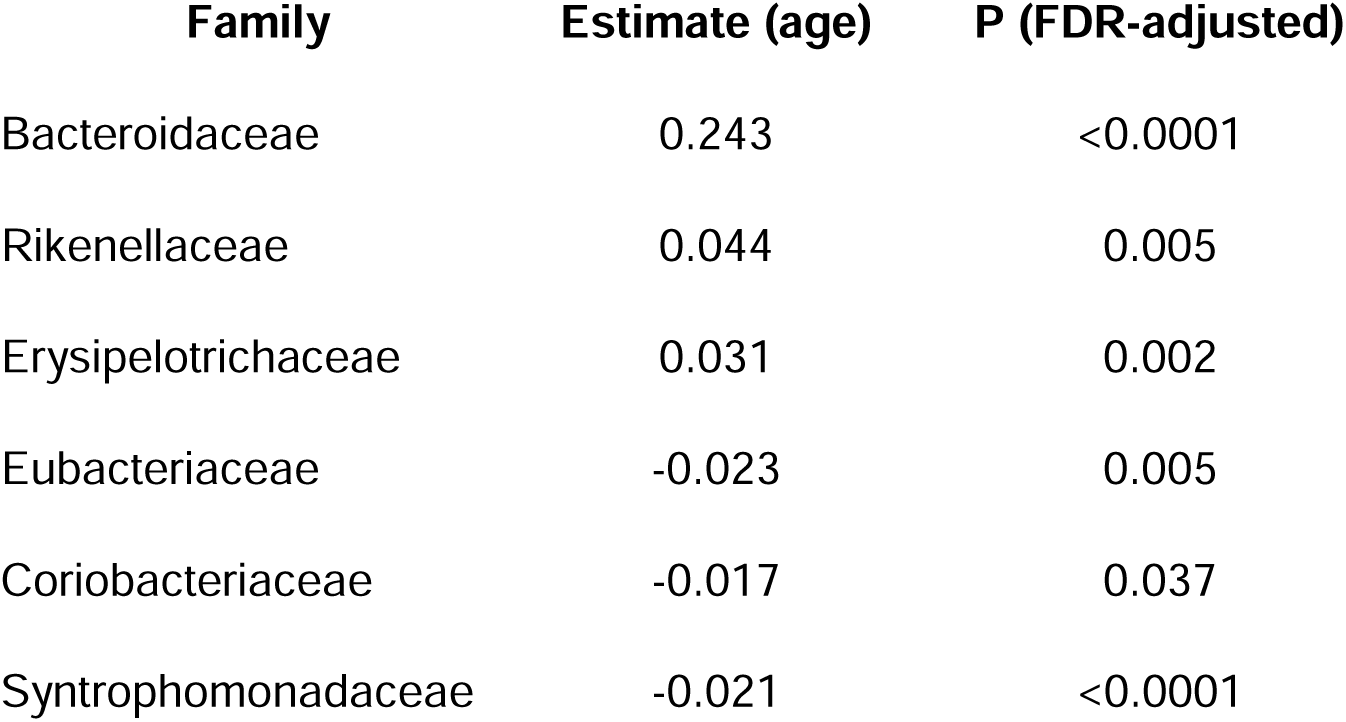
Differentially enriched bacterial families with age class. Results from linear mixed-effects models (dependent variable: arcsine transformed relative abundance) show significant (P_FDR_ ≤ 0.05) differences in the relative abundances of 6 bacterial families. Negative model estimates reflect taxa that are greater in relative abundance in adults; positive estimates reflect taxa that are greater in relative abundance in juveniles. Models controlled for age (adult/juvenile) and sex (male/female) as fixed factors, with date of sample collection and nest box (family membership) as random effects.

**Table S5.**
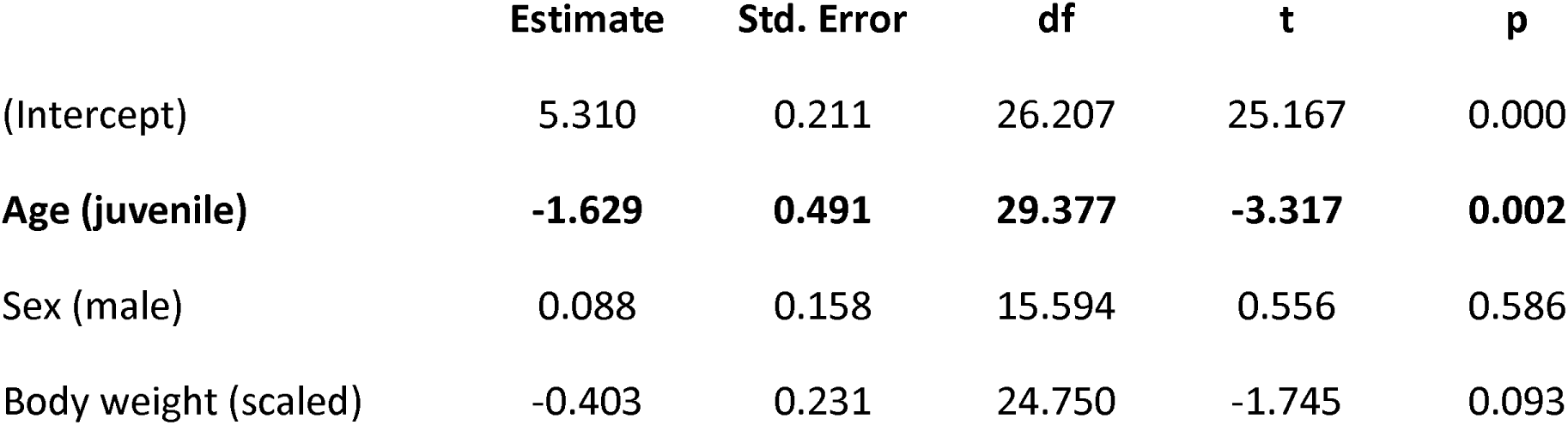
Glucocorticoid production varies with host age. Results from linear mixed-effects model testing for effects of host age and sex on hair GC concentrations (pg/mg, logged), controlling for time of sample collection. Sampling date and family membership as random effects explained no variation and were therefore removed from this and subsequent models in which GCs were the dependent variable to avoid singular fitting of the model.

|  | **Estimate** | **Std. Error** | **df** | **t** | **p** |
| --- | --- | --- | --- | --- | --- |
| (Intercept) | 5.310 | 0.211 | 26.207 | 25.167 | 0.000 |
| **Age (juvenile)** | **-1.629** | **0.491** | **29.377** | **-3.317** | **0.002** |
| Sex (male) | 0.088 | 0.158 | 15.594 | 0.556 | 0.586 |
| Body weight (scaled) | -0.403 | 0.231 | 24.750 | -1.745 | 0.093 |

**Table S6.** Age effects on gut microbial alpha-diversity. Results from linear mixed-effects models testing the effect of host age and sex on gut microbial (A) richness and (B) abundance-weighted diversity. Models controlled for sampling date and family membership as random effects. Shannon diversity was tukey-transformed to achieve residual normality prior to modeling.

| Richness (# of OTUs) |  |  |  |  |  |
| --- | --- | --- | --- | --- | --- |
|  | Estimate | Std. Error | df | t | p |
| (Intercept) | 831.430 | 63.750 | 38.000 | 13.042 | 0.000 |
| **Age (juvenile)** | **-464.850** | **139.250** | **37.120** | **-3.338** | **0.002** |
| Sex (male) | 35.180 | 51.570 | 37.660 | 0.682 | 0.499 |
| Body weight (scaled) | -69.380 | 69.870 | 37.910 | -0.993 | 0.327 |
| Shannon diversity (abundance-weighted) |  |  |  |  |  |
| (Intercept) | 70.550 | 8.656 | 34.976 | 8.150 | 0.000 |
| **Age (juvenile)** | **-55.944** | **18.116** | **36.160** | **-3.088** | **0.004** |
| Sex (male) | -6.093 | 6.710 | 35.661 | -0.908 | 0.370 |
| Body weight (scaled) | -7.783 | 9.122 | 36.308 | -0.853 | 0.399 |

**Table S7.** Covariation between glucocorticoids and gut microbial diversity. Results from linear-mixed effects models testing the effect of hair GCs on gut microbial alpha-diversity in adults and juveniles. Models controlled for date of sample collection and nest box/family membership.

| **A** |  | **Adults: richness (# of OTUs)** |  |  |  |
| --- | --- | --- | --- | --- | --- |
|  | **Estimate** | **Std. Error** | **df** | **t** | **p** |
| (Intercept) | 796.210 | 14.470 | 25.560 | 55.009 | 0.000 |
| Glucocorticoids (pg/mg) | 26.820 | 14.020 | 19.670 | 1.914 | 0.070 |
| **B** |  | **Adults: Shannon diversity (abundance-weighted)** |  |  |  |
| (Intercept) | 63.420 | 5.900 | 3.174 | 10.750 | 0.001 |
| **Glucocorticoids (pg/mg)** | **0.475** | **0.017** | **12.775** | **28.610** | **0.000** |
| **C** |  | **Juveniles: richness (# of OTUs)** |  |  |  |
| (Intercept) | 460.168 | 57.173 | 6.816 | 8.049 | 0.000 |
| Glucocorticoids (pg/mg) | 62.686 | 47.809 | 11.985 | 1.311 | 0.214 |
| **D** |  | **Juveniles: Shannon diversity (abundance-weighted)** |  |  |  |
| (Intercept) | 19.120 | 3.483 | 2.189 | 5.490 | 0.026 |
| Glucocorticoids (pg/mg) | 2.573 | 2.624 | 11.645 | 0.981 | 0.347 |

**Table S8.** Similarity in glucocorticoid concentrations predict more similar gut microbiota among pairs of adult squirrels. Results from *brms* model testing the effects of various pairwise environmental and host factors on Jaccard similarity across pairs of adult squirrels. Significant (where 95% credible intervals do not overlap zero) terms shown in bold.

|  | **Estimate** | **Error** | **l-95% CI** | **u-95% CI** | **Rhat** | **Bulk_ESS** | **Tail_ESS** |
| --- | --- | --- | --- | --- | --- | --- | --- |
| Same sex | -0.02 | 0.03 | -0.08 | 0.03 | 1 | 6781 | 5852 |
| **Spatial distance** | **-0.23** | **0.07** | **-0.36** | **-0.11** | **1** | **6652** | **6070** |
| **Same family** | **0.02** | **0.25** | **-0.48** | **0.48** | **1** | **7005** | **4408** |
| **Glucocorticoid difference** | **-0.36** | **0.12** | **-0.6** | **-0.12** | **1** | **4483** | **4797** |

**Table S9.**
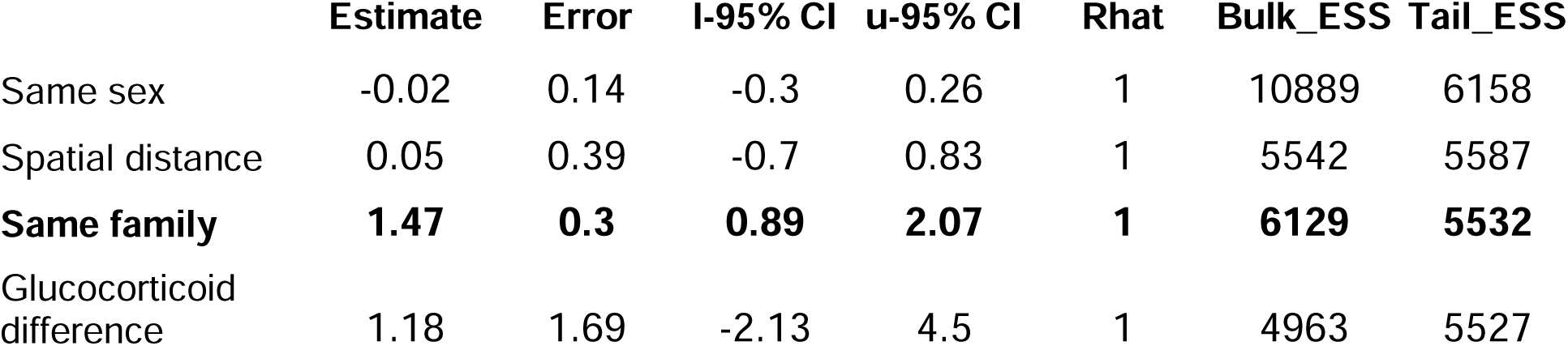
Among pairs of juvenile squirrels, glucocorticoid similarity did not predict microbial similarity. Results from *brms* model testing the effects of various pairwise environmental and host factors on Jaccard similarity across pairs of juvenile squirrels. Significant (where 95% credible intervals do not overlap zero) terms shown in bold.

|  | Estimate | Error | l-95% CI | u-95% CI | Rhat | Bulk_ESS | Tail_ESS |
| --- | --- | --- | --- | --- | --- | --- | --- |
| Same sex | -0.02 | 0.14 | -0.3 | 0.26 | 1 | 10889 | 6158 |
| Spatial distance | 0.05 | 0.39 | -0.7 | 0.83 | 1 | 5542 | 5587 |
| **Same family** | **1.47** | **0.3** | **0.89** | **2.07** | **1** | **6129** | **5532** |
| Glucocorticoid difference | 1.18 | 1.69 | -2.13 | 4.5 | 1 | 4963 | 5527 |

**Table S10.**
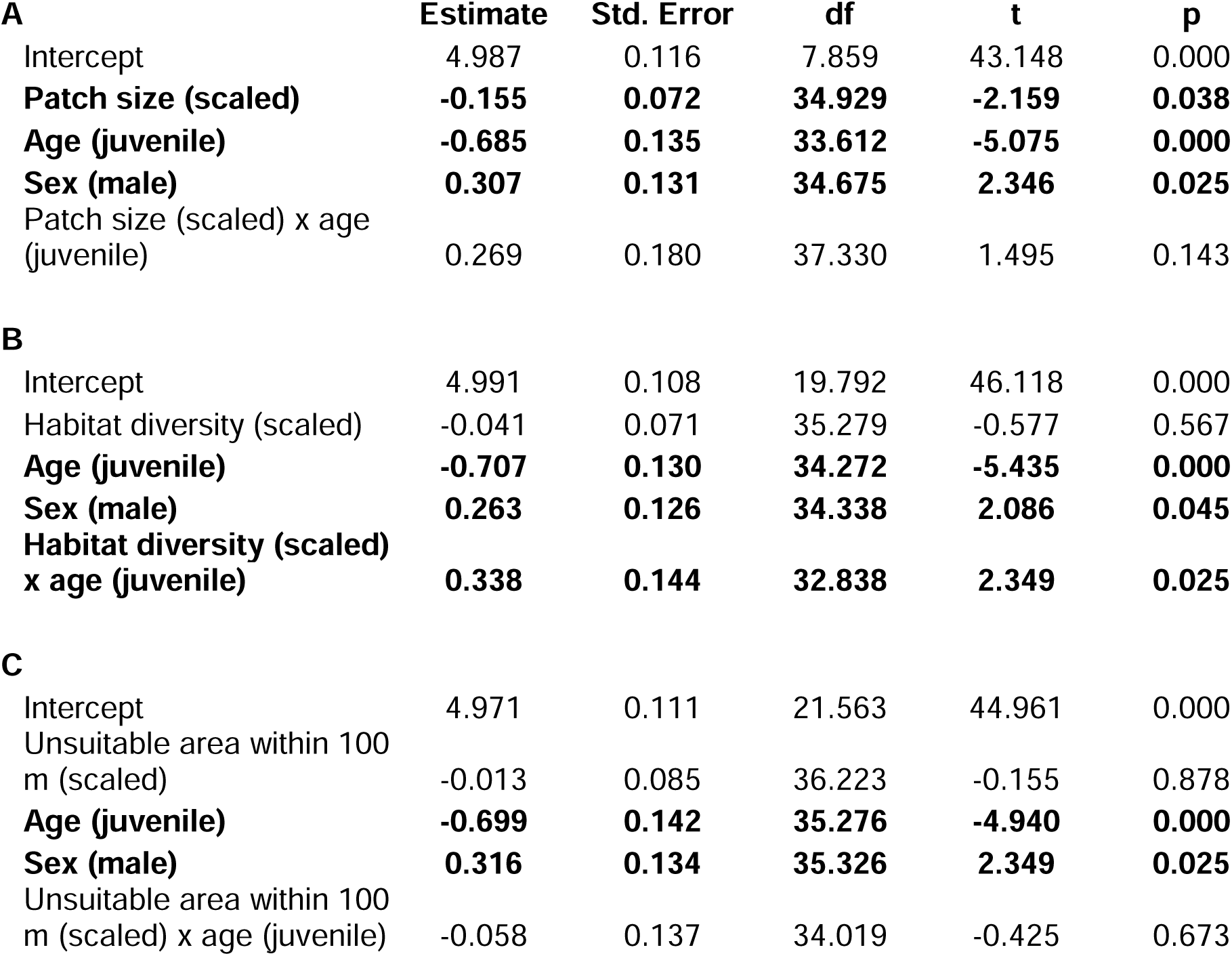
Habitat quality predicts glucocorticoid concentrations in an age-dependent manner. Results show effects of different measures of habitat quality (A: patch size, B: habitat diversity, C: amount of unsuitable area within 100 m) on hair GCs (pg/mg, logged) independently and as a function of age class. Results from linear mixed-effects models controlling for host age and sex (fixed factors) as well as date and time of sample collection (random effects).

**Table S11.**
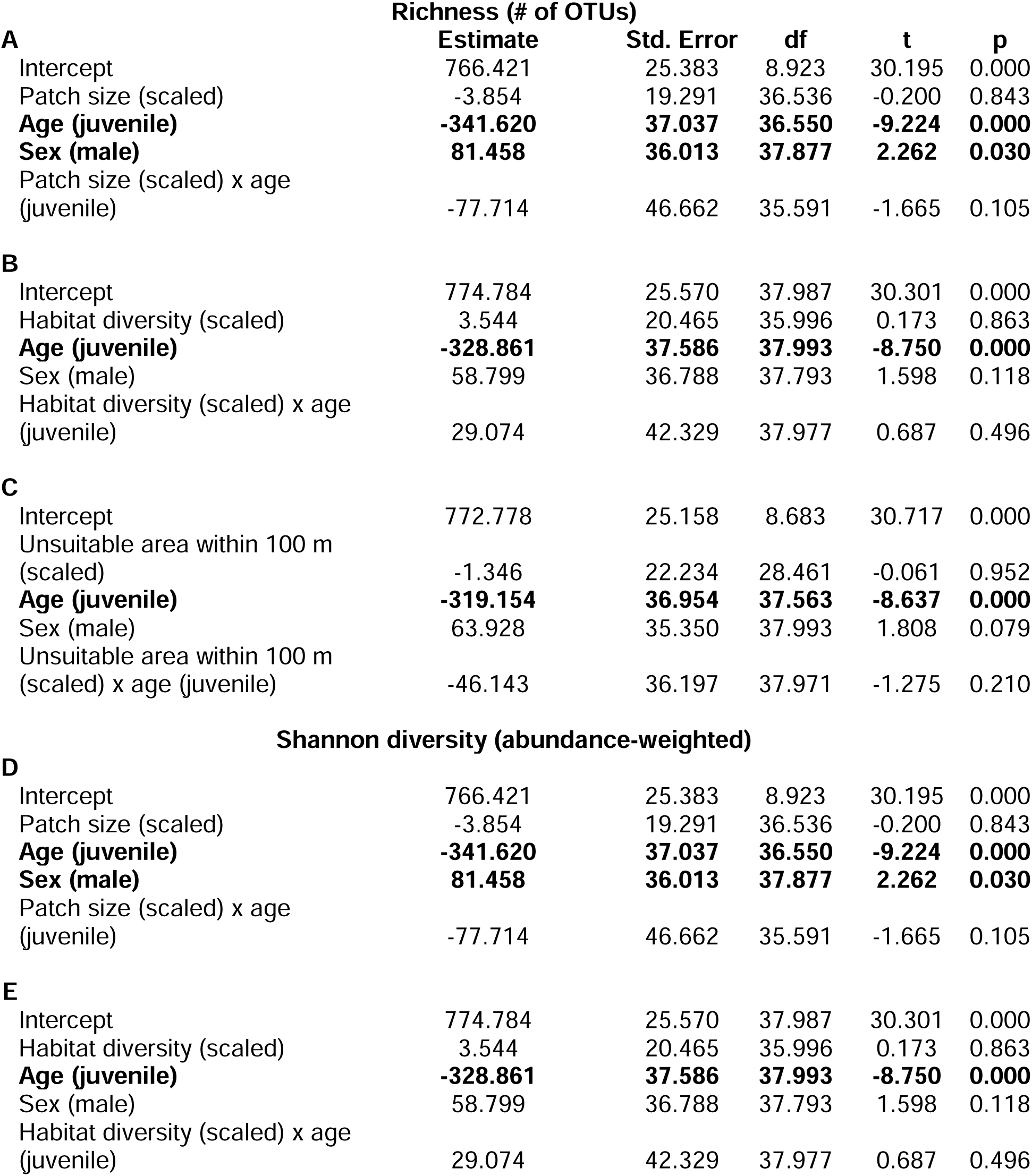
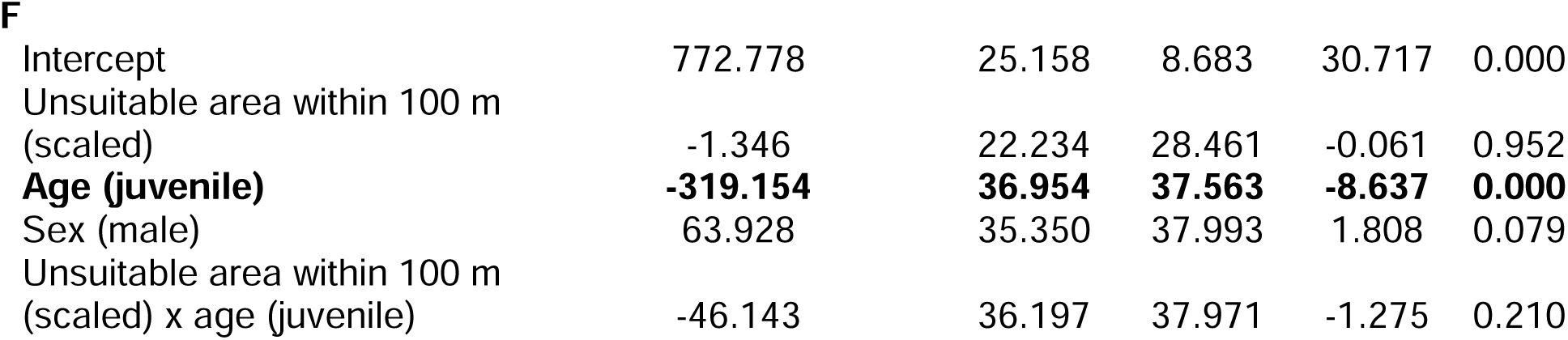
No direct effects of habitat quality on gut microbial richness or abundance-weighted diversity in either age class. Results from generalized linear-mixed effects models testing effects of three habitat variables (patch size, habitat diversity, and amount of unsuitable area within 100 m of an individual’s nest box) on (A) gut microbial richness and (B) Shannon diversity (abundance-weighted alpha-diversity). All models included sampling date and nest box (i.e., family membership) as random effects.

**Table S12.**
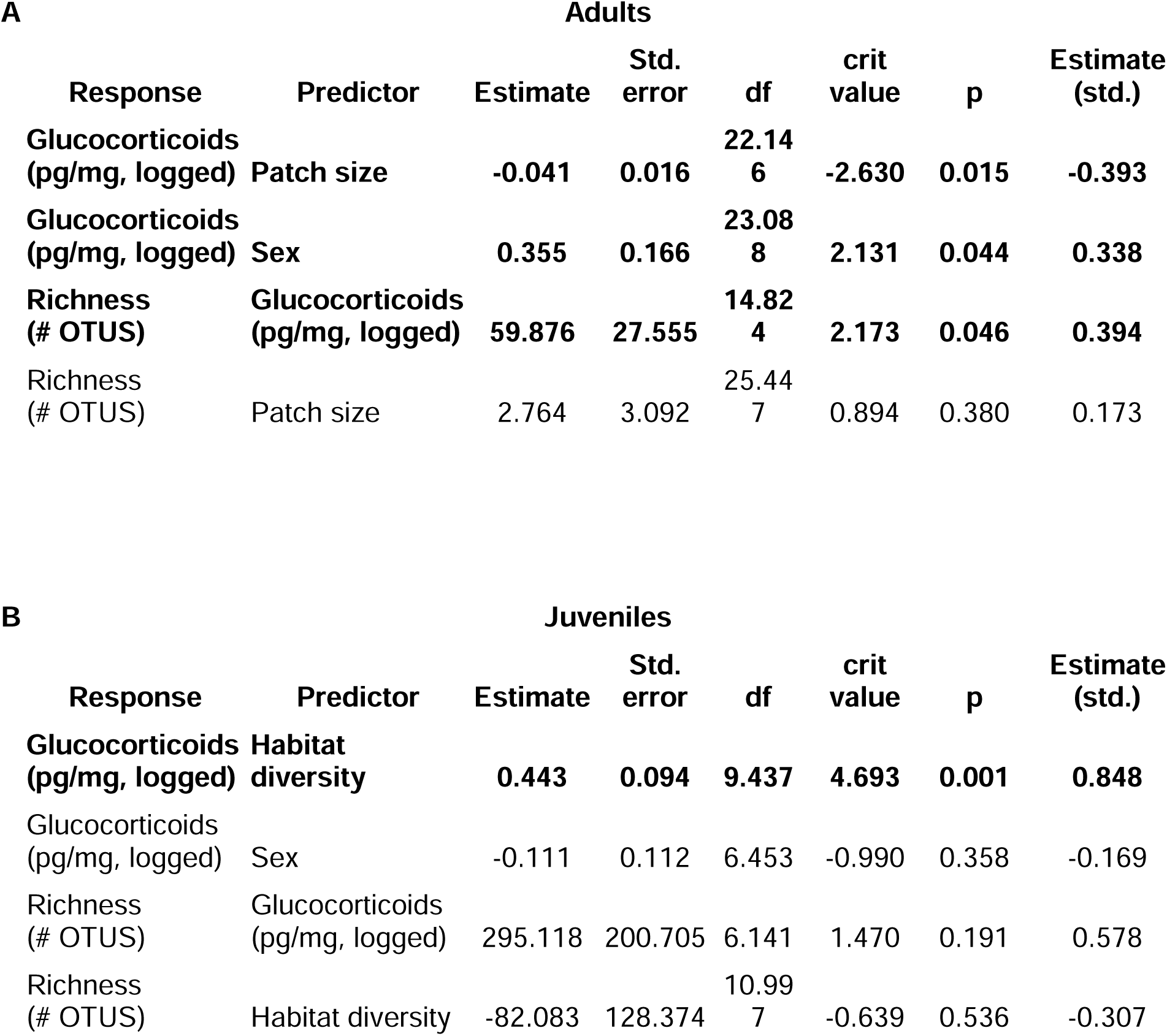
Age-dependent indirect effects of habitat quality on microbial richness. Results from structural equation models testing for direct versus indirect paths connecting habitat quality, hair GCs, and gut microbial richness in adults and juveniles. Models included two components (GCs ∼ habitat quality, and gut microbiota ∼ GCs), controlling for sex, date, time of sample collection and family membership/nest box as random effects. Tests of directional separation indicated that models were not missing any significant paths and fit the data well (adult SEM: Fisher’s C = 0.52, df = 2, *P* = 0.77; juvenile SEM: Fisher’s C = 1.77, df = 2, *P* = 0.41).

## Figures

**Figure S1.**
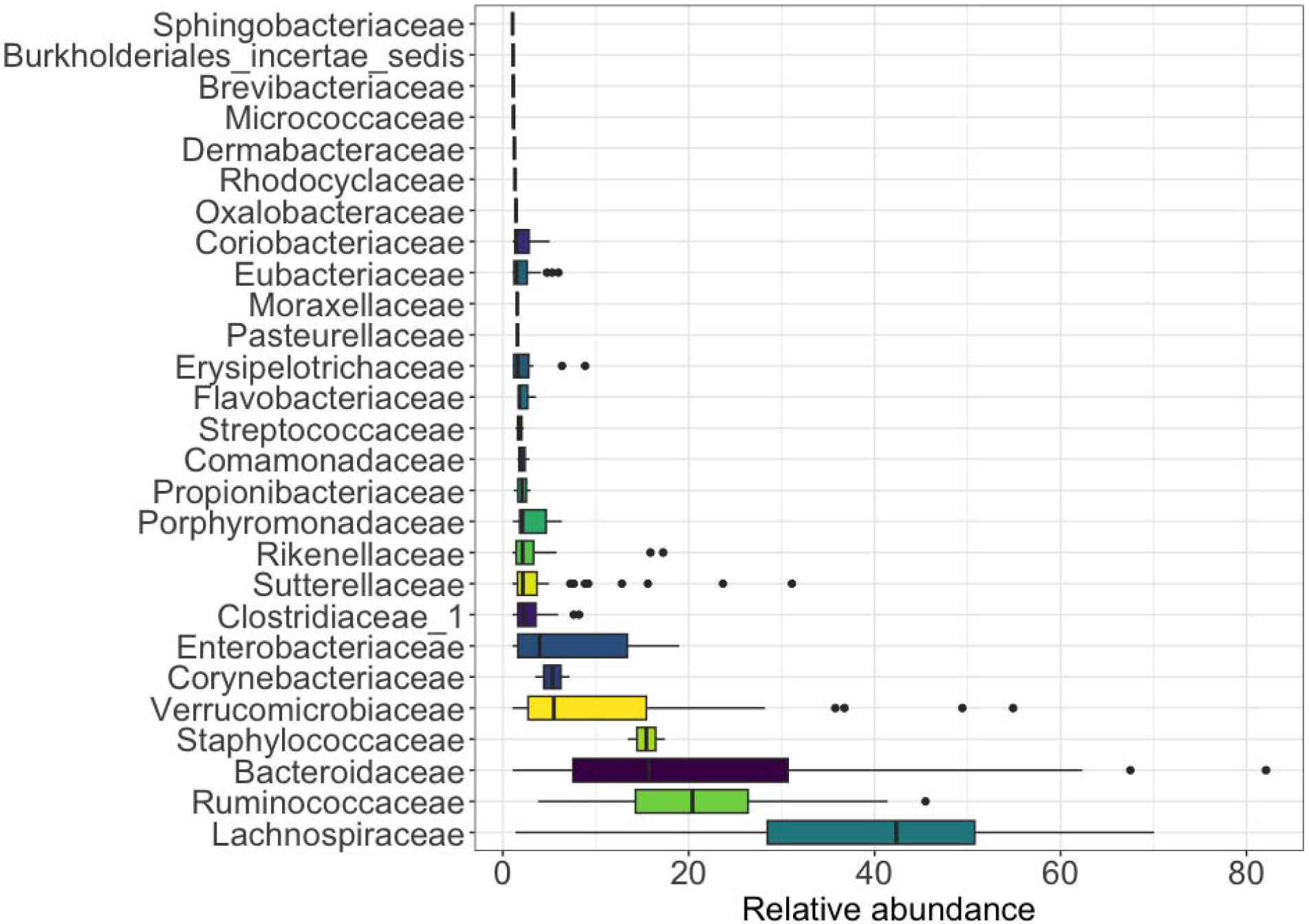
Relative abundance of bacterial families in the Siberian flying squirrel gut microbiota. Families shown reflect the top taxa with at least 5% mean relative abundance across samples.

**Figure S2.**
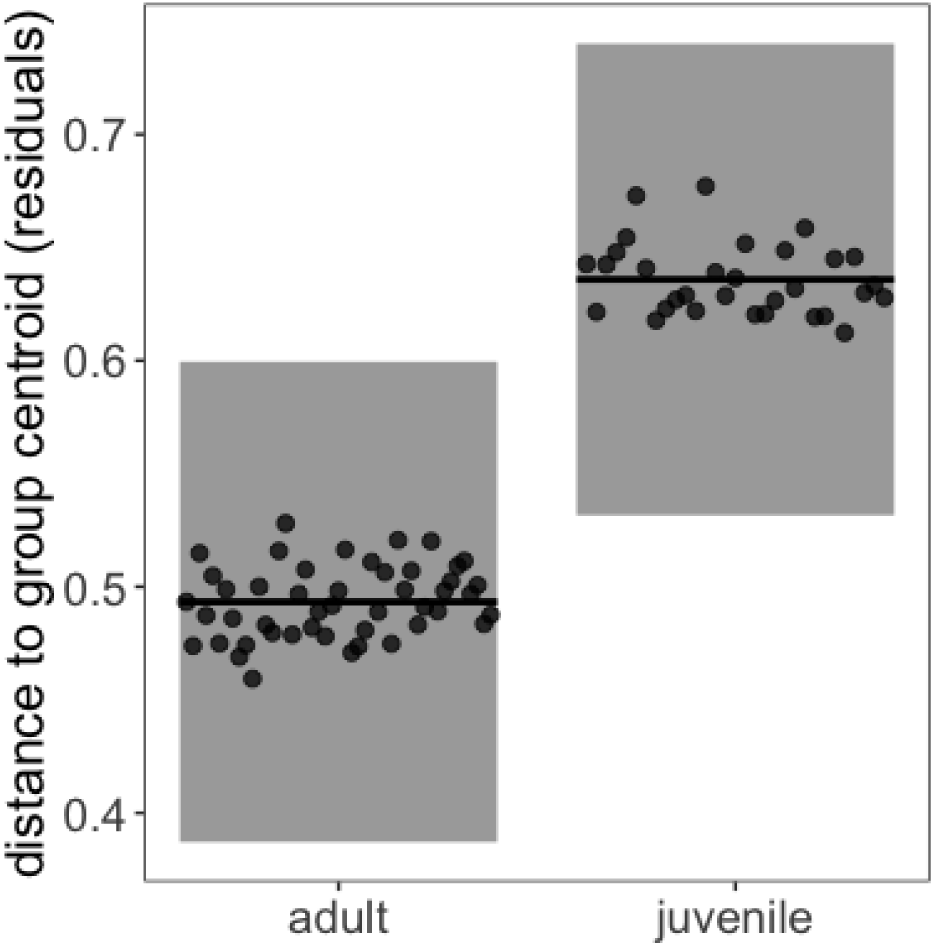
Variation in homogeneity of dispersion suggests differences in gut microbial individuality with age.

**Figure S3.**
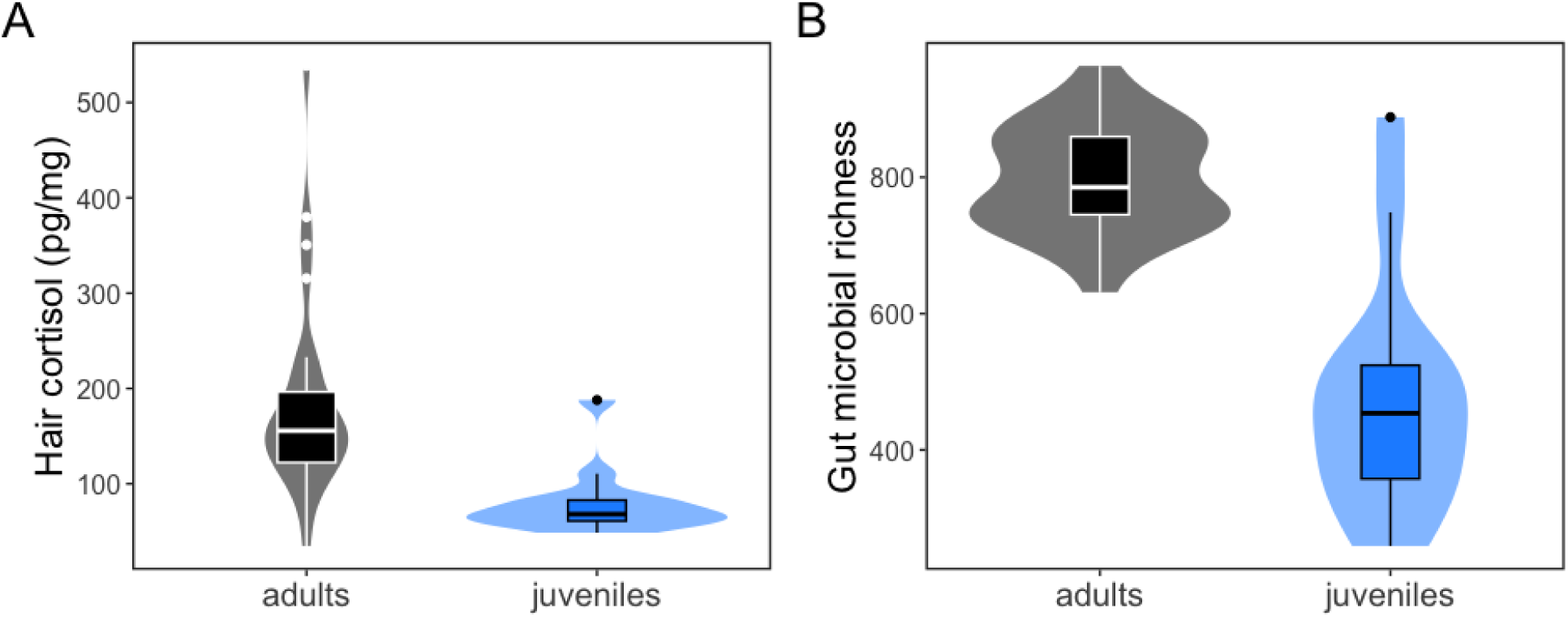
Juveniles exhibit lower hair glucocorticoids and gut microbial richness compared to adults. Box and violin plots depict raw data (lines = median, points = outliers).

## AUTHOR CONTRIBUTIONS

LP carried out the statistical analyses of the data and led writing of the manuscript. AR conceptualised the hypotheses herein, led the data collection, developed statistical methods for analysis of the data and helped writing the manuscript. AS participated in the data collection, derived the habitat diversity measures from GIS data and provided feedback on the manuscript. RW developed the field system, participated in the data collection and provided feedback on the manuscript. AH helped develop the analysis methods and provided feedback on the manuscript.

