## Supplemental Materials for "Indirect environmental effects on the gut-brain axis in a wild mammal"

### "Habitat quality influences gut microbiota via cortisol in Siberian flying squirrels (Pteromys volans)"

**Table S1. Intrinsic predictors of gut microbial variation**. Results from a marginal PERMANOVA testing effects host age, sex, and family membership/nest box on Jaccard similarity, with 999 permutations.

| | Df | SumOfSqs | R2 | F | p |
|---|---|---|---|---|---|
| **Nest box/family** | **54.000** | **23.801** | **0.716** | **1.263** | **0.001** |
| **Age** | **1.000** | **0.714** | **0.021** | **2.044** | **0.001** |
| Sex | 1.000 | 0.336 | 0.010 | 0.962 | 0.606 |
| Residual | 21.000 | 7.329 | 0.220 | | |
| Total | 77.000 | 33.240 | 1.000 | | |

**Table S2. Extrinsic and methodological predictors of gut microbial variation**. Results from a marginal PERMANOVA testing effects of habitat quality (habitat diversity, patch size, and amount of unsuitable area within 100 m of an individual’s nest box) and date of sample collection on gut microbial composition (Jaccard similarity), controlling for nest box/family membership (blocking factor).

| | Df | SumOfSqs | R2 | F | p |
|---|---|---|---|---|---|
| **Age** | **1.000** | **1.340** | **0.066** | **3.432** | **0.001** |
| Habitat diversity | 1.000 | 0.441 | 0.022 | 1.130 | 0.129 |
| Patch size | 1.000 | 0.463 | 0.023 | 1.185 | 0.072 |
| Unsuitable area within 100 m | 1.000 | 0.421 | 0.021 | 1.078 | 0.216 |
| **Sampling date** | **3.000** | **1.540** | **0.075** | **1.315** | **0.002** |
| Residual | 41.000 | 16.013 | 0.785 | | |
| Total | 48.000 | 20.410 | 1.000 | | |

**Table S3. Juvenile flying squirrels exhibit greater inter-individual variability in gut microbial composition than adults.** Results from generalized linear mixed-effects model (family = beta) testing the effects of age class on distance to group centroid in a Jaccard similarity compositional space. Model controlled for sampling date, family membership/nest box, and host sex.

| | **Estimate** | **Std. Error** | **z** | **p** |
|---|---|---|---|---|
| **Intercept** | **0.507** | **0.021** | **24.242** | **0.000** |
| **Age (juvenile)** | **0.143** | **0.026** | **5.401** | **0.000** |
| Sex (male) | -0.013 | 0.025 | -0.528 | 0.597 |

**Table S4. Differentially enriched bacterial families with age class.** Results from linear mixed-effects models (dependent variable: arcsine transformed relative abundance) show significant ($P_{FDR} \leq 0.05$) differences in the relative abundances of 6 bacterial families. Negative model estimates reflect taxa that are greater in relative abundance in adults; positive estimates reflect taxa that are greater in relative abundance in juveniles. Models controlled for age (adult/juvenile) and sex (male/female) as fixed factors, with date of sample collection and nest box (family membership) as random effects.

| **Family** | **Estimate (age)** | **P (FDR-adjusted)** |
|---|---|---|
| Bacteroidaceae | 0.243 | <0.0001 |
| Rikenellaceae | 0.044 | 0.005 |
| Erysipelotrichaceae | 0.031 | 0.002 |
| Eubacteriaceae | -0.023 | 0.005 |
| Coriobacteriaceae | -0.017 | 0.037 |
| Syntrophomonadaceae | -0.021 | <0.0001 |

**Table S5. Cortisol production varies with host age.** Results from linear mixed-effects model testing for effects of host age and sex on hair cortisol concentrations (pg/mg, logged), controlling for time of sample collection. Sampling date and family membership as random effects explained no variation and were therefore removed from this and subsequent models in which cortisol was the dependent variable to avoid singular fitting of the model.

| | **Estimate** | **Std. Error** | **df** | **t** | **p** |
|---|---|---|---|---|---|
| (Intercept) | 4.974 | 0.098 | 37.247 | 50.534 | 0.000 |
| **Age (juvenile)** | **-0.791** | **0.127** | **20.304** | **-6.237** | **0.000** |
| Sex (male) | 0.249 | 0.125 | 28.500 | 1.991 | 0.056 |

**Table S6. Age effects on gut microbial alpha-diversity.** Results from linear mixed-effects models testing the effect of host age and sex on gut microbial (A) richness and (B) abundance-weighted diversity. Models controlled for sampling date and family membership as random effects. Shannon diversity was tukey-transformed to achieve residual normality prior to modeling.

**A** — **Richness (# of OTUs)**

| | **Estimate** | **Std. Error** | **df** | **t** | **p** |
|---|---|---|---|---|---|
| **(Intercept)** | **63.992** | **4.098** | **6.643** | **15.617** | **0.000** |
| **Age (juvenile)** | **-40.865** | **4.758** | **38.081** | **-8.588** | **0.000** |
| Sex (male) | -2.311 | 4.648 | 39.391 | -0.497 | 0.622 |

**B** — **Shannon diversity (abundance-weighted)**

| | | | | | |
|---|---|---|---|---|---|
| **(Intercept)** | **63.992** | **4.098** | **6.643** | **15.617** | **0.000** |
| **Age (juvenile)** | **-40.865** | **4.758** | **38.081** | **-8.588** | **0.000** |
| Sex (male) | -2.311 | 4.648 | 39.391 | -0.497 | 0.622 |

**Table S7. Covariation between cortisol and gut microbial diversity.** Results from linear-mixed effects models testing the effect of hair cortisol concentrations on gut microbial alpha-diversity in adults and juveniles. Models controlled for date of sample collection and nest box/family membership.

| **A** | **Adults: richness (# of OTUs)** | | | | |
|---|---|---|---|---|---|
| | **Estimate** | **Std. Error** | **df** | **t** | **p** |
| (Intercept) | 796.210 | 14.470 | 25.560 | 55.009 | 0.000 |
| Cortisol (pg/mg) | 26.820 | 14.020 | 19.670 | 1.914 | 0.070 |
| **B** | **Adults: Shannon diversity (abundance-weighted)** | | | | |
| (Intercept) | 63.420 | 5.900 | 3.174 | 10.750 | 0.001 |
| **Cortisol (pg/mg)** | **0.475** | **0.017** | **12.775** | **28.610** | **0.000** |
| **C** | **Juveniles: richness (# of OTUs)** | | | | |
| (Intercept) | 460.168 | 57.173 | 6.816 | 8.049 | 0.000 |
| Cortisol (pg/mg) | 62.686 | 47.809 | 11.985 | 1.311 | 0.214 |
| **D** | **Juveniles: Shannon diversity (abundance-weighted)** | | | | |
| (Intercept) | 19.120 | 3.483 | 2.189 | 5.490 | 0.026 |
| Cortisol (pg/mg) | 2.573 | 2.624 | 11.645 | 0.981 | 0.347 |

**Table S8. Similarity in cortisol concentrations predict more similar gut microbiota among pairs of adult squirrels.** Results from *brms* model testing the effects of various pairwise environmental and host factors on Jaccard similarity across pairs of adult squirrels. Significant (where 95% credible intervals do not overlap zero) terms shown in bold.

| | **Estimate** | **Error** | **l-95% CI** | **u-95% CI** | **Rhat** | **Bulk_ESS** | **Tail_ESS** |
|---|---|---|---|---|---|---|---|
| Same sex | -0.02 | 0.03 | -0.08 | 0.03 | 1 | 6781 | 5852 |
| **Spatial distance** | **-0.23** | **0.07** | **-0.36** | **-0.11** | **1** | **6652** | **6070** |
| **Same family** | **0.02** | **0.25** | **-0.48** | **0.48** | **1** | **7005** | **4408** |
| **Cortisol difference** | **-0.36** | **0.12** | **-0.6** | **-0.12** | **1** | **4483** | **4797** |

**Table S9. Among pairs of juvenile squirrels, cortisol similarity did not predict microbial similarity.** Results from *brms* model testing the effects of various pairwise environmental and host factors on Jaccard similarity across pairs of juvenile squirrels. Significant (where 95% credible intervals do not overlap zero) terms shown in bold.

| | Estimate | Error | l-95% CI | u-95% CI | Rhat | Bulk_ESS | Tail_ESS |
|---|---|---|---|---|---|---|---|
| Same sex | -0.02 | 0.14 | -0.3 | 0.26 | 1 | 10889 | 6158 |
| Spatial distance | 0.05 | 0.39 | -0.7 | 0.83 | 1 | 5542 | 5587 |
| **Same family** | **1.47** | **0.3** | **0.89** | **2.07** | **1** | **6129** | **5532** |
| Cortisol difference | 1.18 | 1.69 | -2.13 | 4.5 | 1 | 4963 | 5527 |

**Table S10. Habitat quality predicts cortisol concentrations in an age-dependent manner.** Results show effects of different measures of habitat quality (A: patch size, B: habitat diversity, C: amount of unsuitable area within 100 m) on hair cortisol (pg/mg, logged) independently and as a function of age class. Results from linear mixed-effects models controlling for host age and sex (fixed factors) as well as date and time of sample collection (random effects).

| **A** | **Estimate** | **Std. Error** | **df** | **t** | **p** |
|---|---|---|---|---|---|
| Intercept | 4.987 | 0.116 | 7.859 | 43.148 | 0.000 |
| **Patch size (scaled)** | **-0.155** | **0.072** | **34.929** | **-2.159** | **0.038** |
| **Age (juvenile)** | **-0.685** | **0.135** | **33.612** | **-5.075** | **0.000** |
| **Sex (male)** | **0.307** | **0.131** | **34.675** | **2.346** | **0.025** |
| Patch size (scaled) x age (juvenile) | 0.269 | 0.180 | 37.330 | 1.495 | 0.143 |
| **B** | | | | | |
| Intercept | 4.991 | 0.108 | 19.792 | 46.118 | 0.000 |
| Habitat diversity (scaled) | -0.041 | 0.071 | 35.279 | -0.577 | 0.567 |
| **Age (juvenile)** | **-0.707** | **0.130** | **34.272** | **-5.435** | **0.000** |
| **Sex (male)** | **0.263** | **0.126** | **34.338** | **2.086** | **0.045** |
| **Habitat diversity (scaled) x age (juvenile)** | **0.338** | **0.144** | **32.838** | **2.349** | **0.025** |
| **C** | | | | | |
| Intercept | 4.971 | 0.111 | 21.563 | 44.961 | 0.000 |
| Unsuitable area within 100 m (scaled) | -0.013 | 0.085 | 36.223 | -0.155 | 0.878 |
| **Age (juvenile)** | **-0.699** | **0.142** | **35.276** | **-4.940** | **0.000** |
| **Sex (male)** | **0.316** | **0.134** | **35.326** | **2.349** | **0.025** |
| Unsuitable area within 100 m (scaled) x age (juvenile) | -0.058 | 0.137 | 34.019 | -0.425 | 0.673 |

**Table S11. No direct effects of habitat quality on gut microbial richness or abundance-weighted diversity in either age class.** Results from generalized linear-mixed effects models testing effects of three habitat variables (patch size, habitat diversity, and amount of unsuitable area within 100 m of an individual's nest box) on (A) gut microbial richness and (B) Shannon diversity (abundance-weighted alpha-diversity). All models included sampling date and nest box (i.e., family membership) as random effects.

| | Richness (# of OTUs) | | | | |
|---|---|---|---|---|---|
| **A** | **Estimate** | **Std. Error** | **df** | **t** | **p** |
| Intercept | 766.421 | 25.383 | 8.923 | 30.195 | 0.000 |
| Patch size (scaled) | -3.854 | 19.291 | 36.536 | -0.200 | 0.843 |
| **Age (juvenile)** | **-341.620** | **37.037** | **36.550** | **-9.224** | **0.000** |
| **Sex (male)** | **81.458** | **36.013** | **37.877** | **2.262** | **0.030** |
| Patch size (scaled) x age (juvenile) | -77.714 | 46.662 | 35.591 | -1.665 | 0.105 |
| **B** | | | | | |
| Intercept | 774.784 | 25.570 | 37.987 | 30.301 | 0.000 |
| Habitat diversity (scaled) | 3.544 | 20.465 | 35.996 | 0.173 | 0.863 |
| **Age (juvenile)** | **-328.861** | **37.586** | **37.993** | **-8.750** | **0.000** |
| Sex (male) | 58.799 | 36.788 | 37.793 | 1.598 | 0.118 |
| Habitat diversity (scaled) x age (juvenile) | 29.074 | 42.329 | 37.977 | 0.687 | 0.496 |
| **C** | | | | | |
| Intercept | 772.778 | 25.158 | 8.683 | 30.717 | 0.000 |
| Unsuitable area within 100 m (scaled) | -1.346 | 22.234 | 28.461 | -0.061 | 0.952 |
| **Age (juvenile)** | **-319.154** | **36.954** | **37.563** | **-8.637** | **0.000** |
| Sex (male) | 63.928 | 35.350 | 37.993 | 1.808 | 0.079 |
| Unsuitable area within 100 m (scaled) x age (juvenile) | -46.143 | 36.197 | 37.971 | -1.275 | 0.210 |
| | **Shannon diversity (abundance-weighted)** | | | | |
| **D** | | | | | |
| Intercept | 766.421 | 25.383 | 8.923 | 30.195 | 0.000 |
| Patch size (scaled) | -3.854 | 19.291 | 36.536 | -0.200 | 0.843 |
| **Age (juvenile)** | **-341.620** | **37.037** | **36.550** | **-9.224** | **0.000** |
| **Sex (male)** | **81.458** | **36.013** | **37.877** | **2.262** | **0.030** |
| Patch size (scaled) x age (juvenile) | -77.714 | 46.662 | 35.591 | -1.665 | 0.105 |
| **E** | | | | | |
| Intercept | 774.784 | 25.570 | 37.987 | 30.301 | 0.000 |
| Habitat diversity (scaled) | 3.544 | 20.465 | 35.996 | 0.173 | 0.863 |
| **Age (juvenile)** | **-328.861** | **37.586** | **37.993** | **-8.750** | **0.000** |
| Sex (male) | 58.799 | 36.788 | 37.793 | 1.598 | 0.118 |
| Habitat diversity (scaled) x age (juvenile) | 29.074 | 42.329 | 37.977 | 0.687 | 0.496 |

| | | | | | |
|---|---|---|---|---|---|
| **F** | | | | | |
| Intercept | 772.778 | 25.158 | 8.683 | 30.717 | 0.000 |
| Unsuitable area within 100 m (scaled) | -1.346 | 22.234 | 28.461 | -0.061 | 0.952 |
| **Age (juvenile)** | **-319.154** | **36.954** | **37.563** | **-8.637** | **0.000** |
| Sex (male) | 63.928 | 35.350 | 37.993 | 1.808 | 0.079 |
| Unsuitable area within 100 m (scaled) x age (juvenile) | -46.143 | 36.197 | 37.971 | -1.275 | 0.210 |

**Table S12. Age-dependent indirect effects of habitat quality on microbial richness.** Results from structural equation models testing for direct versus indirect paths connecting habitat quality, hair cortisol, and gut microbial richness in adults and juveniles. Models included two components (cortisol ~ habitat quality, and gut microbiota ~ cortisol), controlling for sex and date and time of sample collection and family membership/nest box as random effects.

**A** **Adults**

| Response | Predictor | Estimate | Std. error | df | crit value | p | Estimate (std.) |
|---|---|---|---|---|---|---|---|
| **Cortisol (pg/mg, logged)** | **Patch size** | **-0.041** | **0.016** | **22.146** | **-2.630** | **0.015** | **-0.393** |
| **Cortisol (pg/mg, logged)** | **Sex** | **0.355** | **0.166** | **23.088** | **2.131** | **0.044** | **0.338** |
| **Richness (# OTUS)** | **Cortisol (pg/mg, logged)** | **59.876** | **27.555** | **14.824** | **2.173** | **0.046** | **0.394** |
| Richness (# OTUS) | Patch size | 2.764 | 3.092 | 25.447 | 0.894 | 0.380 | 0.173 |

**B** **Juveniles**

| Response | Predictor | Estimate | Std. error | df | crit value | p | Estimate (std.) |
|---|---|---|---|---|---|---|---|
| **Cortisol (pg/mg, logged)** | **Habitat diversity** | **0.443** | **0.094** | **9.437** | **4.693** | **0.001** | **0.848** |
| Cortisol (pg/mg, logged) | Sex | -0.111 | 0.112 | 6.453 | -0.990 | 0.358 | -0.169 |
| Richness (# OTUS) | Cortisol (pg/mg, logged) | 295.118 | 200.705 | 6.141 | 1.470 | 0.191 | 0.578 |
| Richness (# OTUS) | Habitat diversity | -82.083 | 128.374 | 10.997 | -0.639 | 0.536 | -0.307 |

**Figures**

**Figure S1. Relative abundance of bacterial families in the Siberian flying squirrel gut microbiota.** Families shown reflect the top taxa with at least 5% mean relative abundance across samples.

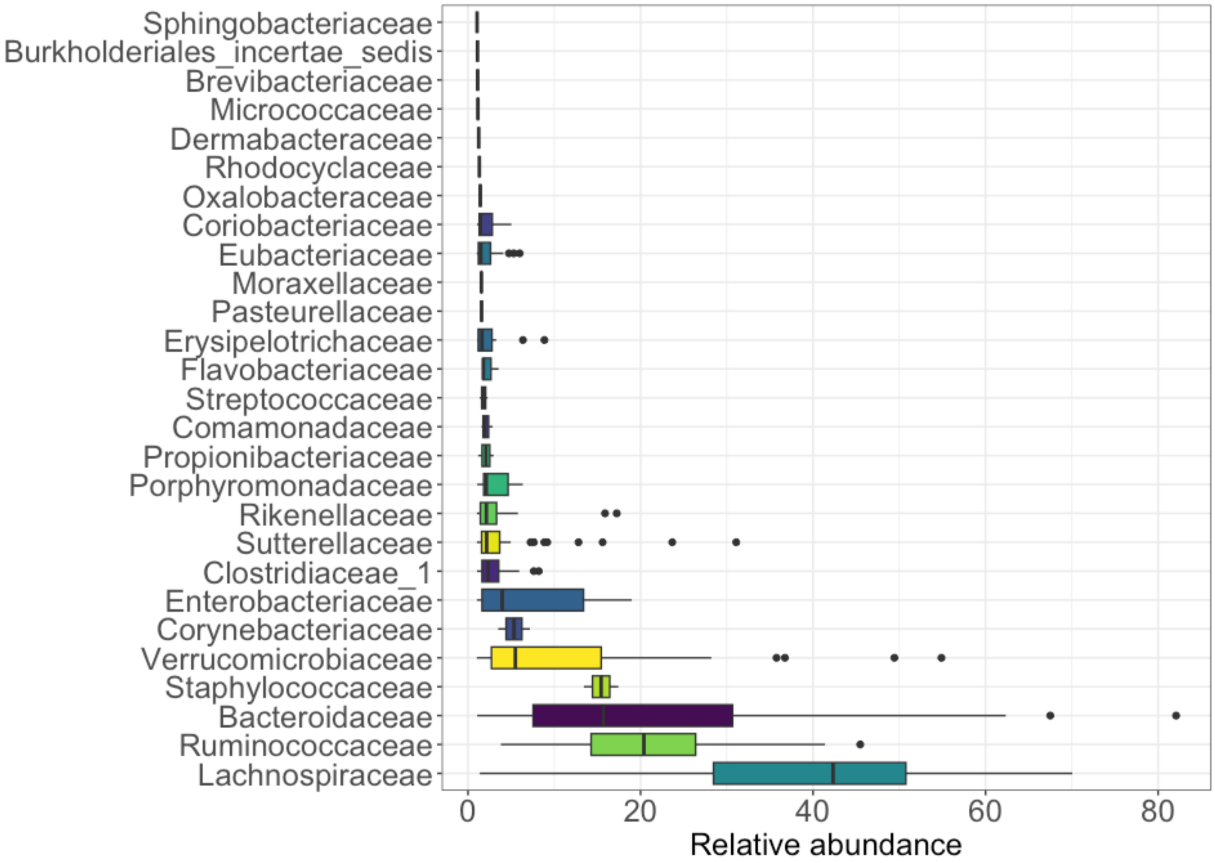

**Figure S2. Variation in homogeneity of dispersion suggests differences in gut microbial individuality with age.**

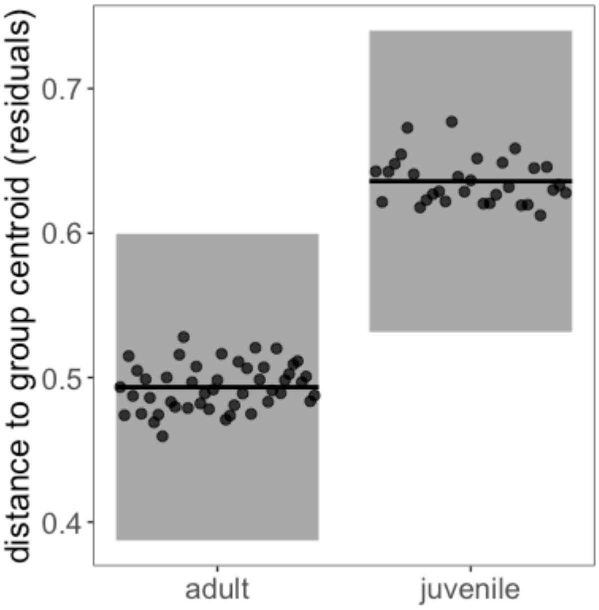

**Figure S3. Juveniles exhibit lower hair cortisol and gut microbial richness compared to adults.** Box and violin plots depict raw data (lines = median, points = outliers).

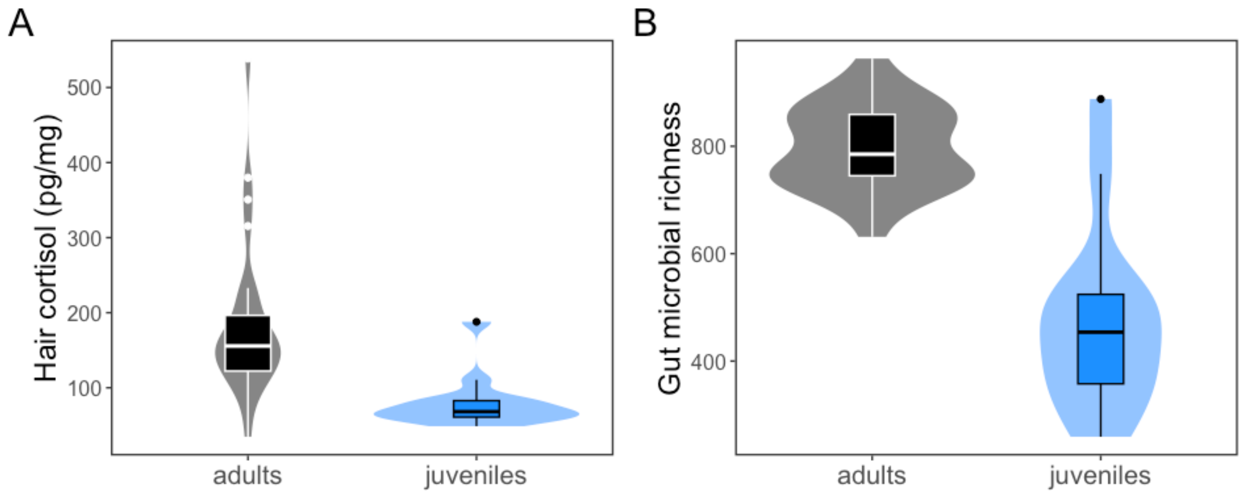
